# Geometric–Chemical Distance Between Protein Surfaces

**DOI:** 10.64898/2026.03.10.710722

**Authors:** Himanshu Swami, John M. McBride, Jean-Pierre Eckmann, Tsvi Tlusty

## Abstract

Proteins recognize, bind, and catalyze through molecular surfaces, where geometry and chemical patterning determine interaction. Comparing these surfaces requires both a geometric–chemical distance and a correspondence that relates one complete surface to another.

Here we introduce IFACE (Intrinsic Field–Aligned Coupled Embedding). IFACE derives a symmetric geometric–chemical distance by optimizing a probabilistic coupling over intrinsic geometry, mean curvature, electrostatics, hydrophobicity, and hydrogen-bond propensity. The same coupling provides an explicit surface map.

For molecular-dynamics conformers, IFACE distinguishes the same protein from distinct proteins more accurately than TM-distance and a Laplace–Beltrami spectral distance. A Jensen–Shannon distribution distance performs best in this binary identity test, because aggregate surface-feature distributions already identify each protein.

A distance must also satisfy a global requirement: its pairwise values must place many distinct protein surfaces consistently in one space. We therefore tested IFACE across six protein families. It produces the strongest family classification and clustering among the distributional, spectral, MaSIF, and SurfaceID comparisons.

The inferred maps preserve geodesic neighborhoods and transfer heme-centered pocket regions across cytochrome P450 proteins. IFACE therefore provides, from one construction, both a distance between complete protein surfaces and the local map that explains that distance.

## INTRODUCTION

Molecular recognition, catalysis, regulation, and assembly occur at protein surfaces^1–6^. More broadly, protein form and function are shaped by the interplay among mechanics, chemistry and evolution^7–12^. The relationship between sequence and folded structure is well established^13,14^, and modern sequence-based methods now predict folded structures with high accuracy^15,16^. Recent work further indicates that these models can capture subtle mutation-induced structural responses^17–20^. Yet a central problem remains: how to compare proteins at the level of their *interfaces*^21–23^. Functional specificity often resides at the exposed surface rather than in the global fold^24,25^.

A protein surface is a curved interface formed by solvent-exposed side chains^26^, where geometry and chemistry jointly determine interaction. Curvature constrains approach; electrostatics, hydrophobicity, and hydrogen-bonding shape interaction specificity^27–31^. These chemical features are themselves statistically correlated with surface curvature^32^. The molecular boundary is not unique: common choices include the van der Waals surface, the solvent-accessible surface (SAS)^33^, and the solvent-excluded surface (SES)^34^. Here we use SES, which approximates the interface encountered by solvent and binding partners; the framework is not tied to this particular surface definition.

Despite the physical coupling of geometry and chemistry at protein surfaces, existing approaches typically compare surfaces through global shape, spectral geometry, descriptors, or learned local representations. In contrast, we formulate protein-surface comparison as a correspondence problem in which intrinsic geometry, extrinsic curvature, and spatially distributed chemical fields jointly determine the map, and a symmetric distance between the complete surfaces is derived from that map.

Protein-surface comparison has developed along two complementary lines. Classical methods compare shape, intrinsic geometry, or physicochemical descriptors, often through global summaries or surface projections^35–42^. Learning-based methods, including MaSIF, DeeplyTough, DeepSurf, and SurfaceID, integrate geometric and physicochemical information in representations designed for interface recognition, binding-site detection, and local patch retrieval^43–47^. These approaches provide powerful measures of surface similarity, but do not generally yield both a coherent correspondence between complete surfaces and a symmetric distance derived from that correspondence.

Correspondence-based methods in shape analysis instead construct mappings between objects, ranging from rigid and non-rigid registration to functional maps and related spectral formulations^48–55^. Protein surfaces, however, are rough, often non-isometric, and may contain cavities and narrow channels, making stable geometric landmarks difficult to define. More importantly, interaction specificity depends on chemical patterning as well as shape^56^. The missing object is therefore a correspondence between complete protein surfaces determined jointly by intrinsic geometry and distributed physicochemical fields, together with a symmetric distance derived from that correspondence.

Conceptually, a protein surface is a curved manifold with intrinsic geometric distances and spatially organized physicochemical fields. A meaningful notion of similarity therefore relates fields across the underlying geometry. This requires a correspondence between surfaces: without one, geometric distortion and physicochemical mismatch cannot be combined into a single distance.

Here we introduce IFACE (Intrinsic Field–Aligned Coupled Embedding), which realizes this construction through a probabilistic coupling that provides bidirectional soft correspondences and defines a symmetric geometric–chemical distance between complete protein surfaces.

We demonstrate the framework in two complementary settings. First, we test whether the method distinguishes conformers of the same protein from different proteins. In this benchmark, IFACE combines intrinsic geometric distortion with mismatches in mean curvature, hydrogen-bond propensity, hydrophobicity, and electrostatic potential. It separates same-protein conformers from distinct proteins more accurately than TM-distance (1 − TM-score)^57^ and a Laplace–Beltrami spectral distance. A correspondence-free Jensen–Shannon distribution distance performs best in this binary identity test, showing that aggregate surface-feature distributions can already encode protein identity.

Second, we examine similarity at the family level. Here the task is not merely to distinguish same from different: the full set of pairwise distances must organize many distinct protein surfaces according to their mutual relations. Across six protein families, IFACE gives the strongest classification and clustering among the Jensen–Shannon distribution, Laplace–Beltrami spectral, MaSIF, and SurfaceID distances examined. The benchmark includes cytochrome P450 proteins, whose shared fold encompasses substantial functional and surface diversity^58,59^.

Finally, we evaluate the correspondence itself. IFACE preserves local geodesic neighborhoods and transfers heme-centered pocket regions between cytochrome P450 surfaces. Thus, the same variational construction provides both a geometric–chemical distance between complete protein surfaces and an explicit local map that reveals the surface regions underlying that distance.

### The IFACE Method

We introduce IFACE (Intrinsic Field– Aligned Coupled Embedding), a correspondence-based framework for comparing protein surfaces via coupled geometry and chemical fields. Figure 1 summarizes the construction. Given two protein surfaces, *S*_*α*_ and *S*_*β*_, each is represented by its intrinsic geometric structure together with spatially distributed surface fields: mean curvature, electrostatic potential, hydrogen-bond propensity, and hydrophobicity (Fig. 1a). Mean curvature is treated as a field because it is a local scalar measure of extrinsic geometry, distinct from the intrinsic geodesic structure of the surface. We then compute an optimal coupling matrix that aligns the two surfaces by balancing structural consistency with feature-field agreement (Fig. 1b). The relative contribution of these terms is controlled by the parameter *λ*, defined in the variational formulation below.

**FIG. 1.**
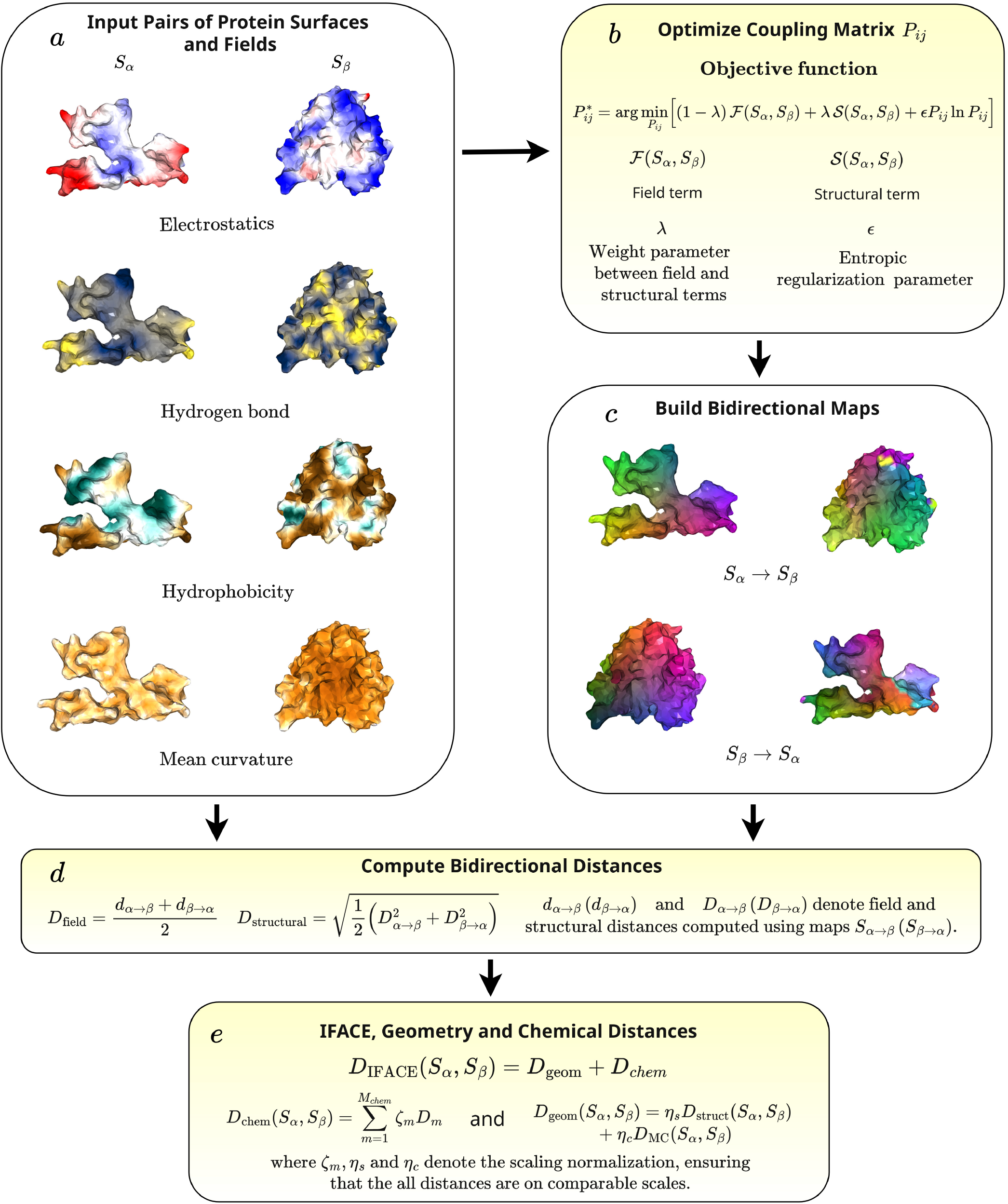
Conceptual workflow of IFACE. (a) Protein surfaces *S*_*α*_ and *S*_*β*_ are represented using geometry and surface feature fields (electrostatics, hydrogen-bond propensity, hydrophobicity, and curvature). (b) An optimal coupling matrix *P*_*i j*_ is computed by balancing structural and feature-field similarity. (c) The coupling defines bidirectional soft correspondences between surfaces. (d) Correspondence-based structural and feature-field distances are computed. (e) Distances are normalized to ensure they are all on the same scale; their combination defines the IFACE distance, which comprises a geometric distance (structural and mean-curvature [MC] components) and a chemical distance (hydrogen-bond, hydrophobicity, and electrostatic components).

The resulting coupling defines soft correspondences between surface points and induces bidirectional surface maps between the two proteins (Fig. 1c). Using these correspondences, we compute symmetric structural and feature-field discrepancies (Fig. 1d). From these quantities, we construct a geometric distance combining intrinsic structural distortion with mean-curvature mismatch, and a chemical distance combining hydrogen-bond propensity, hydrophobicity, and electrostatic mismatch. After normalization to a common scale, their combination defines the geometric– chemical IFACE distance. This procedure yields a symmetric measure that integrates intrinsic and extrinsic geometry with aligned chemical fields while retaining the explicit surface correspondence. Full details of the optimization and definitions are provided in the Methods.

Our implementation depends on a few parameter choices, most notably *λ* and *ϵ* (Fig. 1b). The parameter *λ* controls the balance between intrinsic geometry and surface-field agreement, whereas *ϵ* controls the strength of entropic regularization during optimization. Across broad ranges of both parameters, the principal results and relative performance of the distances remain stable (see SI).

## RESULTS

We evaluate IFACE in three stages: discrimination of molecular-dynamics conformers, organization of proteins across six families, and validation of the inferred surface correspondences. Unless stated otherwise, all surfaces are represented by meshes with 3,000 vertices; details of surface generation are provided in the Supplementary Information. All distances are computed directly from the variational correspondence, without training or learned embeddings.

We begin with a basic but essential consistency test. If the representation faithfully encodes surface organization, distinct conformers of the same protein should remain close to one another, while remaining clearly separated from unrelated proteins, under the IFACE distance. We therefore compare IFACE distances among conformers of the same protein with distances between different proteins. This experiment probes whether conformational variability is correctly distinguished from genuine inter-protein dissimilarity.

We then consider a more demanding regime: organization at the protein-family level across six families. Here, we test whether proteins belonging to the same family group together under the IFACE distance. This setting asks whether the complete matrix of pairwise distances recovers shared family-level surface organization across diverse proteins. Successful classification and clustering indicate that the representation captures conserved geometric–chemical patterns rather than relying on any single global descriptor.

Finally, we evaluate the inferred surface correspondences directly. We test whether mapped vertices preserve local geodesic neighborhoods across the conformer and family benchmarks, and whether the maps transfer heme-centered pocket regions between cytochrome P450 proteins. These analyses assess whether the correspondence underlying the IFACE distance is locally coherent and identifies biologically corresponding surface regions.

### IFACE discerns conformers of the same protein as closer than different proteins

In this task, we use the ATLAS dataset^60^ and focus on four proteins with PDB codes^61^ 6XRX^62^, 5HZ7^63^, 2XZ3^64^, and 6XDS^65^, corresponding to an AEG12 mosquito protein, a DNA-binding protein, a viral protein, and a signaling protein, respectively. These structures are fusion constructs in which the proteins of interest are fused to maltose-binding protein (MBP), a commonly used solubility and crystallization tag. Consequently, each contains the same MBP domain fused to a different protein of interest. All structures were restricted to chain A. ATLAS contains molecular dynamics (MD) trajectories for 1,390 proteins from the Protein Data Bank (PDB), with three independent simulations per protein and 10,001 snapshots per trajectory. Each trajectory spans 100 ns. On this timescale, the native fold is preserved; what fluctuates is the surface. The resulting ensembles therefore sample thermal surface variability without undergoing large conformational rearrangements. This makes ATLAS suitable for assessing whether a distance captures intrinsic surface fluctuations rather than gross structural change.

To construct a stringent benchmark, we searched for proteins that allow controlled comparison both within and between proteins. Sequence analysis using MMseqs2^66^ showed that most ATLAS entries are mutually unrelated, aside from occasional duplicates. A notable exception is a set of four related synthetic constructs fused to maltose-binding protein for crystallization: 6XRX_A^62^, 5HZ7_A^63^, 2XZ3_A^64^, and 6XDS_A^65^. These proteins are similar enough to make discrimination nontrivial, yet distinct enough to test whether combining surface chemistry with geometry adds resolving power beyond global fold similarity.

For each protein, we select ten conformers sampled along the MD trajectory. The resulting TM-score distributions overlap both within and between proteins, creating an ensemble in which intra-protein variability competes directly with inter-protein similarity. This overlap makes the benchmark demanding: it asks whether a surface-based optimal coupling, sensitive to both geometry and physicochemical fields, can resolve distinctions that fold-level similarity cannot.

### Example of IFACE mapping

Before turning to quantitative analysis, we first examine a representative example to visualize the surface correspondence identified by the IFACE optimal transport coupling (Fig. 2). The mapping integrates geometric alignment with multiple surface fields— hydrophobicity, hydrogen-bond propensity, electrostatic potential, and mean curvature—as described above.

**FIG. 2.**
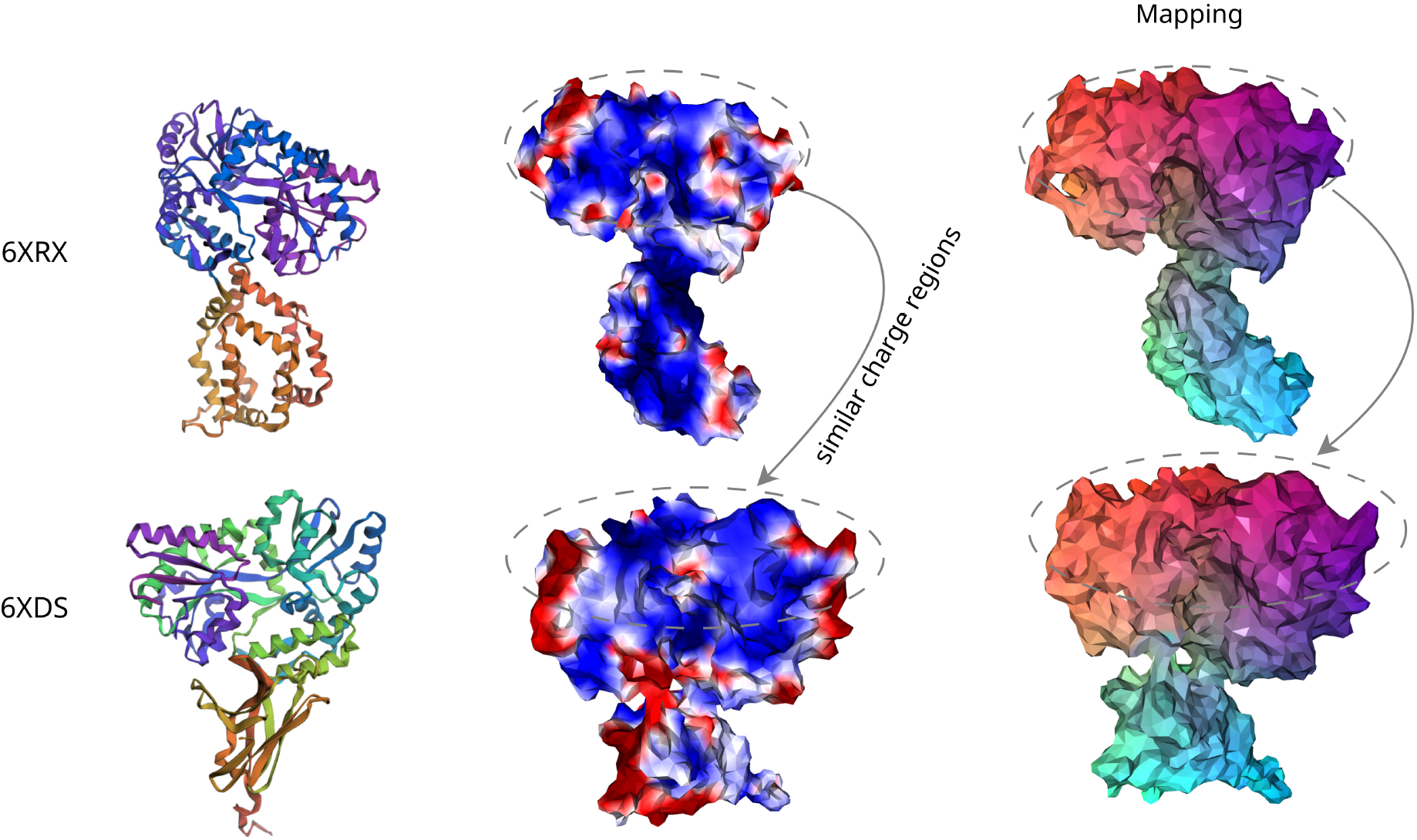
Comparison and mapping of protein surfaces. Protein 6XRX is shown in the top row and 6XDS in the bottom row. Left: ribbon representations of the protein structures. Middle: electrostatic potential distributions (color scale clipped to the 5th–95th percentile of the combined distribution), where similar colors indicate regions with comparable electrostatic potential. Right: IFACE color mappings showing correspondences from 6XRX to 6XDS. The correspondences (vertex mapping) are computed using combined structural and feature-field similarity, allowing direct comparison of corresponding surface regions.

The two proteins shown (6XRX and 6XDS) have a TM-distance of ~0.30 (TM-score ~0.70), indicating substantial global fold similarity. This similarity arises because both constructs contain the same fused maltose-binding protein (MBP) domain, which is identical in sequence in the two structures. The shared domain accounts for a substantial fraction of each construct and therefore produces strong global alignment. The non-MBP proteins have distinct biological functions, so the informative differences lie primarily outside the shared domain, in the non-MBP regions and their exposed surfaces.

The middle column of Fig. 2 displays the electrostatic potential mapped onto the two surfaces. The upper regions of 6XRX (lipid-binding protein) and 6XDS (signaling protein) exhibit closely matched electrostatic profiles and compatible geometry. The IFACE correspondence (rightmost panel) aligns these regions across the two proteins, identifying surface patches that are jointly similar in shape and physicochemical character. In contrast, the lower regions differ in both curvature and electrostatics, and consequently contribute more strongly to the overall distance.

This example illustrates how the optimized surface correspondence integrates geometric and chemical information to resolve surface-level similarity beyond fold alignment alone. Having established this qualitative intuition, we now proceed to a systematic evaluation of whether IFACE reliably separates conformers of the same protein from distinct proteins.

### IFACE distance robustly separates conformers from distinct proteins

We quantify discriminatory power using the *IFACE distance*, a composite measure that integrates geometric and physicochemical features of protein surfaces. Each component—structural (struct), mean curvature (meancurv), hydrophobicity (hphob), hydrogen-bond propensity (hbond), and electrostatic potential (elect)—is normalized using bounds obtained from a fixed reference set, placing the components on comparable numerical scales.

The IFACE distance is then defined as

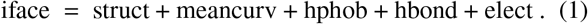

This assigns equal weight to intrinsic structure, mean curvature, and each physicochemical field. This equal-weight choice avoids parameter tuning and provides a transparent baseline against which more elaborate weighting schemes could be assessed (Eq. 1 corresponds to Eq. 4 in Methods with equal weights). For comparison, we also define a *chemical-feature distance*,

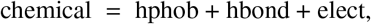

which isolates the contribution of physicochemical fields independent of structural alignment. We also define a *geometry-only distance*,

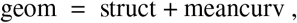

which isolates the contribution of geometry from that of the chemical fields.

For this task, we compare IFACE against three classes of baselines: (i) TM-distance, defined as 1 − TM-score, where TM-score is a widely used structural similarity measure based on structural alignment; (ii) a correspondence-free distributional baseline defined as the average Jensen– Shannon divergence^67^ computed independently for each geometric and physicochemical surface feature; and (iii) an intrinsic spectral distance based on the Laplace–Beltrami (LB) operator^68^. These baselines provide complementary comparisons based on global structural similarity, feature distributions, and intrinsic geometry, respectively.

Figure 3 summarizes the binary classification performance and feature ablation analysis on the conformer benchmark. For the componentwise ablation comparison, the four scalar surface fields are matched directly between IFACE and JSD. Because the intrinsic structural component of IFACE has no direct distributional counterpart, it is compared with the JSD of the Gaussian-curvature distribution as an additional geometric descriptor. The precision–recall (Fig. 3a) and ROC (Fig. 3b) curves show that both IFACE and the distribution-based JSD representation accurately distinguish same-protein conformers from distinct proteins, outperforming the Laplace–Beltrami (LB) distance and the TM-score baseline. For this benchmark, JSD achieves the highest overall performance (AP = 0.984 vs. 0.928; ROC-AUC = 0.994 vs. 0.950), indicating that aggregate descriptor distributions are especially effective in this binary proteinidentity test.

**FIG. 3.**
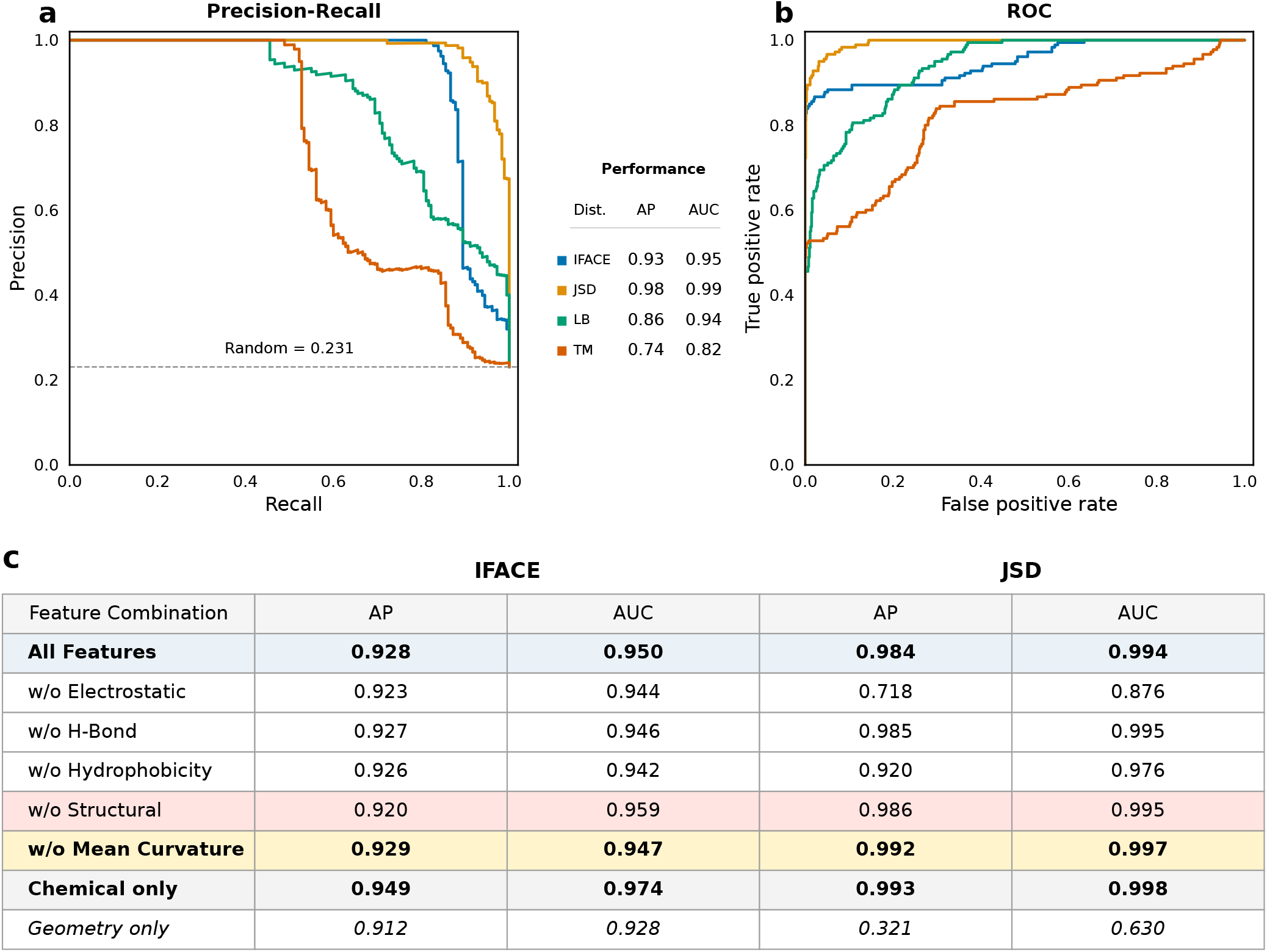
Conformer benchmark classification and feature ablation. (a) Precision–Recall curves and (b) Receiver–Operating Characteristic (ROC) curves comparing IFACE, JSD, Laplace–Beltrami (LB), and TM distances. Performance is summarized by Average Precision (AP) and ROC–AUC. IFACE outperforms both TM and LB, while JSD achieves the highest classification performance on this benchmark. (c) Leave-one-feature-out ablation comparing correspondence-based IFACE with matched distribution-based JSD distances. Removing individual descriptors has only a modest effect on IFACE, indicating that its performance is robust to individual feature removal. In contrast, JSD exhibits greater sensitivity to descriptor selection: removing the electrostatic feature substantially reduces performance, while the geometry-only representation performs markedly worse than its correspondence-based IFACE counterpart.

The feature contribution analysis (Fig. 3c) provides further insight into the role of individual descriptors. Electrostatics is the dominant descriptor for JSD: removing it reduces AP from 0.984 to 0.718 and ROC-AUC from 0.994 to 0.876. The corresponding reduction for IFACE is only modest, from AP = 0.928 to 0.923 and ROC-AUC = 0.950 to 0.944. Removing the remaining descriptors likewise produces only minor variations in IFACE performance, whereas JSD varies more strongly for Electrostatics.

Chemical descriptors alone slightly outperform the full five-feature representation for both methods (IFACE: AP = 0.949, ROC-AUC = 0.974; JSD: AP = 0.993, ROC-AUC = 0.998). By contrast, geometry-only JSD performs substantially worse (AP = 0.321, ROC-AUC = 0.630), whereas geometry-only IFACE shows only a small degradation. Thus, JSD discrimination in this benchmark depends strongly on physicochemical information, particularly electrostatics, whereas no single descriptor is solely responsible for the performance of IFACE.

### Family-level similarity of proteins

In this experiment, we evaluate whether the proposed distance measures organize proteins coherently at the family level. Unlike the binary conformer benchmark, this task tests whether the full matrix of pairwise distances recovers relationships among many distinct protein surfaces. This framework yields a distance-based representation of protein space and enables direct comparisons across protein families without relying on sequence or fold annotations.

To evaluate family-level protein surface similarity, we constructed a benchmark comprising 25 proteins in total: 24 from six structurally and functionally distinct protein families and one unrelated outlier (PDB ID: 3L40) (Table I). The benchmark includes eight cytochrome P450 proteins, three histone proteins, three serine proteases, four globins, two C-type lysozymes, and four members of the ribonuclease A superfamily, as well as a single unrelated outlier. These families were selected because they exhibit substantial diversity in biological function, sequence, and structural organization, while remaining sufficiently well characterized to provide representative examples of each family. The proteins originate from evolutionarily diverse organisms, including bacteria, a bacteriophage, an amoeba, humans, cattle, pigs, sperm whales, chickens, frogs, fruit flies, and fission yeast. This selection introduces considerable evolutionary diversity both within and across protein families.

**TABLE I.** Proteins used in the family-level clustering benchmark, their source organisms, and common organism classifications.

| PDB ID | Protein family/group | Source organism | Common name/type |
| --- | --- | --- | --- |
| 2IJ5 | Cytochrome P450 | *Mycobacterium tuberculosis* | Bacterium |
| 6N6Q | Cytochrome P450 | Mycobacterium phage Adler | Bacteriophage |
| 3OFU | Cytochrome P450 | *Novosphingobium aromaticivorans* | Bacterium |
| 6ZI2 | Cytochrome P450 | *Streptomyces antibioticus* | Bacterium |
| 6UX0 | Cytochrome P450 | *Acanthamoeba castellanii* | Amoeba |
| 4H23 | Cytochrome P450 | *Priestia megaterium* | Bacterium |
| 1JPZ | Cytochrome P450 | *Priestia megaterium* | Bacterium |
| 1TQN | Cytochrome P450 | *Homo sapiens* | Human |
| 1AOI | Histone proteins | *Xenopus laevis* | African clawed frog |
| 5Y0C | Histone proteins | *Homo sapiens* | Human |
| 4QLC | Histone proteins | *Drosophila melanogaster*; *Gallus gallus* | Fruit fly; chicken |
| 1BTP | Serine proteases | *Bos taurus* | Cattle |
| 1ESA | Serine proteases | *Sus scrofa* | Pig |
| 1GCD | Serine proteases | *Bos taurus* | Cattle |
| 2DN1 | Globins | *Homo sapiens* | Human |
| 1A6M | Globins | *Physeter macrocephalus* | Sperm whale |
| 1OJ6 | Globins | *Homo sapiens* | Human |
| 1UT0 | Globins | *Homo sapiens* | Human |
| 1LYZ | C-type lysozymes | *Gallus gallus* | Chicken |
| 1LZ1 | C-type lysozymes | *Homo sapiens* | Human |
| 7RSA | Ribonuclease A superfamily | *Bos taurus* | Cattle |
| 1ANG | Ribonuclease A superfamily | *Homo sapiens* | Human |
| 1GQV | Ribonuclease A superfamily | *Homo sapiens* | Human |
| 2RNF | Ribonuclease A superfamily | *Homo sapiens* | Human |
| 3L40 | Outlier protein | *Schizosaccharomyces pombe* | Fission yeast |

Our benchmark is designed to determine whether surface-based similarity measures recover biologically meaningful family relationships rather than merely reflecting sequence identity or global structural resemblance. Proteins belonging to the same family are expected to cluster together despite evolutionary divergence, whereas proteins from different families should remain well separated. The inclusion of a structurally unrelated outlier further assesses the robustness of the distance measures when applied to proteins that do not naturally belong to any benchmark family. Together, these features make the set a stringent benchmark for family-level classification, clustering, and correspondence-based surface comparison.

### Example of pocket mapping within the same protein family

We illustrate surface correspondence within the P450 family using full-length human cytochrome P450 3A4 (PDB ID: 1TQN, sequence length 486, chain A) and P450 BM-3 from *Bacillus megaterium* (PDB ID: 1JPZ, sequence length 473, chain A). Both proteins contain a heme prosthetic group coordinated by a conserved cysteine residue. The heme pocket is deeply buried, with substrate access occurring through channels or tunnels. Its internal topology—comprising cavities, handles, and narrow passages—renders it difficult to identify using surface-comparison schemes such as conformal mapping or projection onto a plane or sphere.

Figure 4 compares proteins 1JPZ and 1TQN. The top row shows ribbon representations of 1JPZ (left) and 1TQN (right), with the heme prosthetic group highlighted in red. The bottom-left image highlights the surface patch surrounding the heme in 1JPZ in red. This buried patch is accessed through substrate tunnels and surrounds the catalytic heme, where substrate oxidation occurs through oxygen activation, exemplifying nontrivial internal surface geometry. The bottom-right image shows the corresponding mapped region in 1TQN (blue), identifying the homologous buried pocket.

**FIG. 4.**
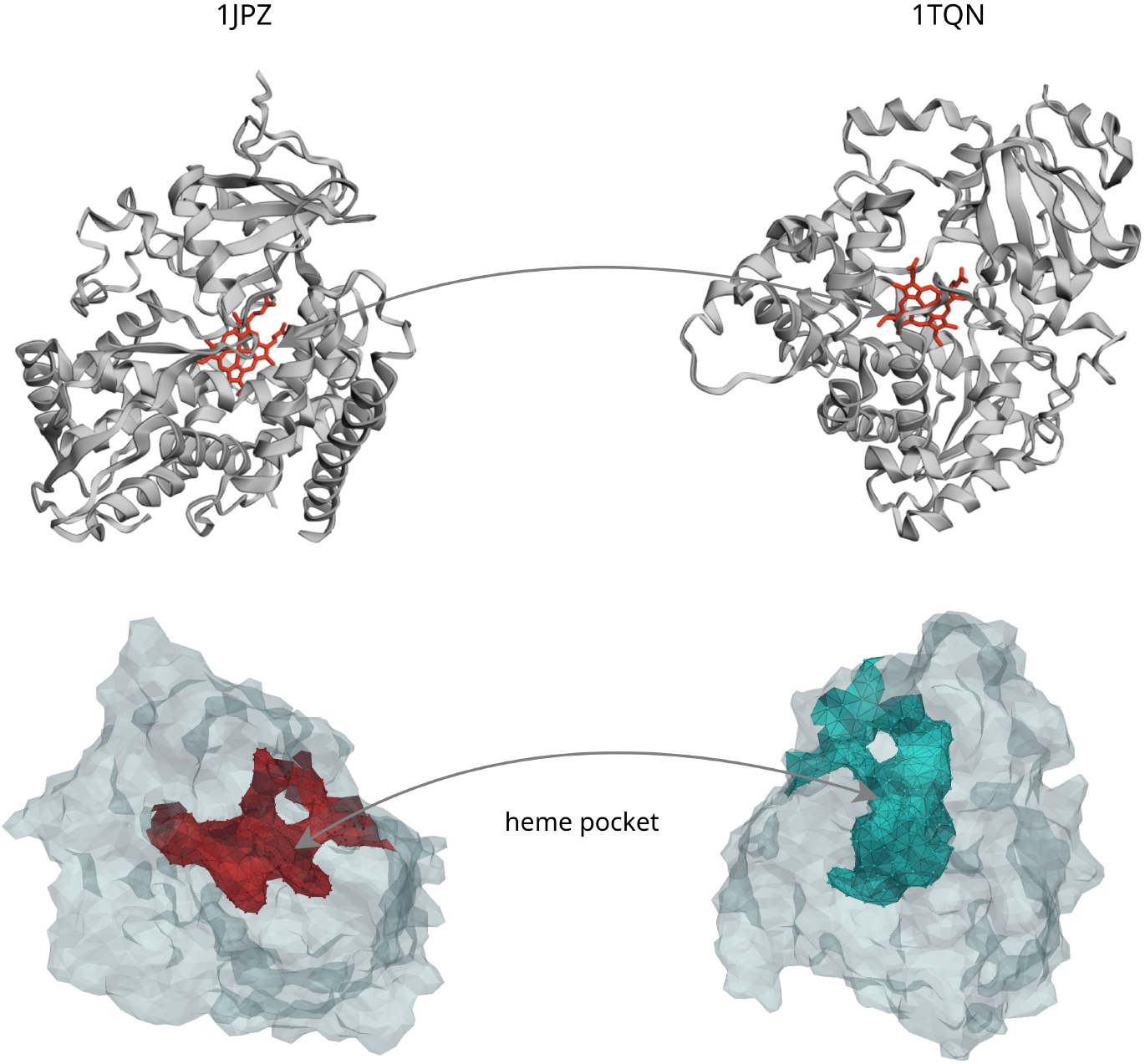
Comparison of proteins 1JPZ and 1TQN. Top row: ribbon representations of chain A from 1JPZ (left) and 1TQN (right), with the heme prosthetic group shown in red. Bottom left: patch around the heme on 1JPZ (red), containing the heme group responsible for substrate oxidation via oxygen activation. This patch lies within the protein interior and is accessed through substrate tunnels, illustrating non-trivial internal surface geometry. Bottom right: the region on 1TQN obtained by mapping the heme-centered pocket of 1JPZ with IFACE, illustrating correspondence between the buried pocket regions of the two proteins.

The probabilistic surface correspondence recovers this conserved internal organization despite substantial geometric complexity. The alignment identifies corresponding heme-centered regions without requiring global topological simplification, indicating that the coupling captures intrinsic surface organization rather than superficial shape similarity.

### Surface-distance-based clustering of protein surfaces

In this task, using the family benchmark, we compare protein pairs from the same family (intra-family) with pairs drawn from different families (inter-family). As shown in Fig. 5(a), we formulate the comparison as a binary classification task, with intra-family pairs as positives and inter-family pairs as negatives.

**FIG. 5.**
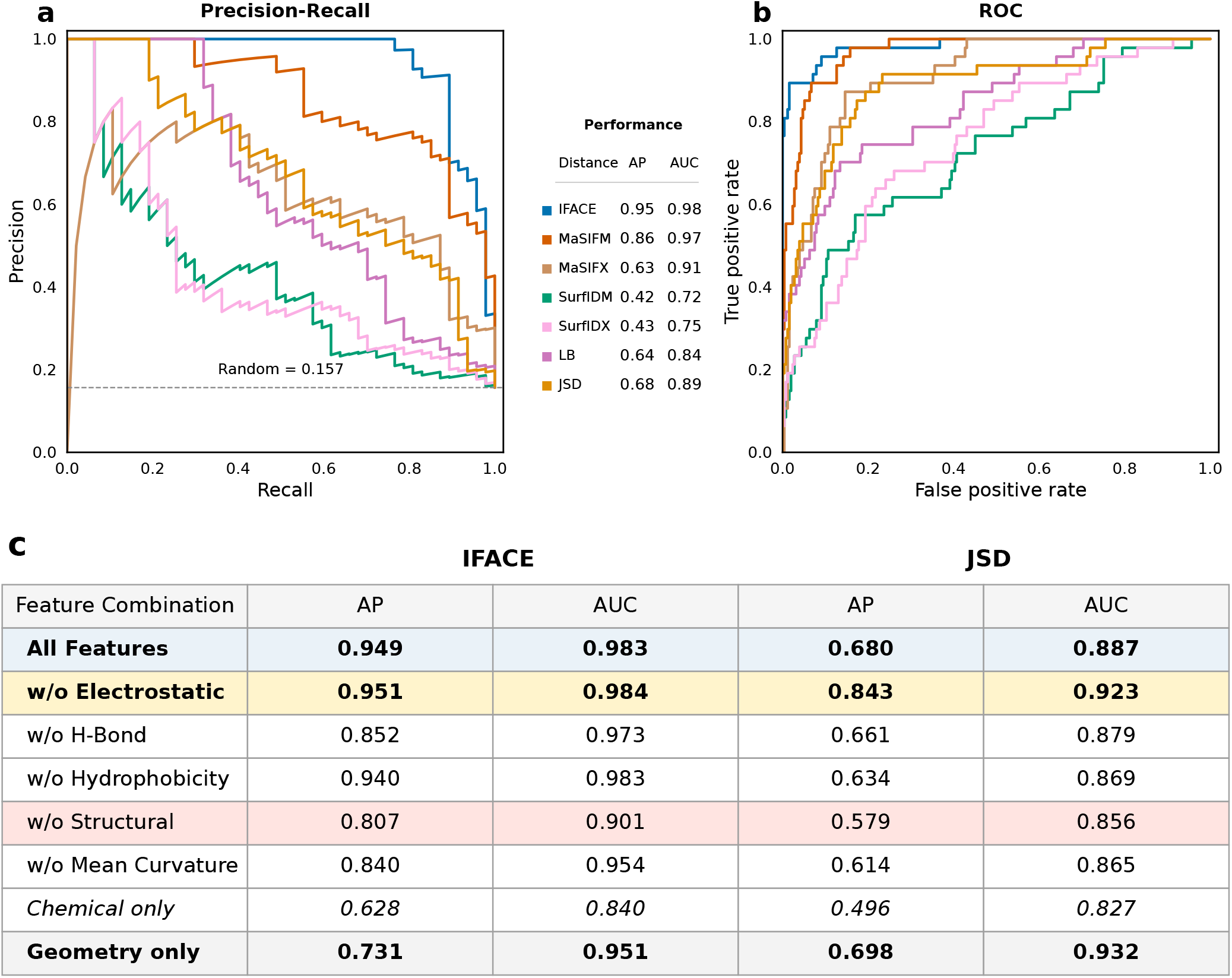
Family benchmark classification and feature ablation. (a) Precision–Recall curves and (b) Receiver–Operating Characteristic (ROC) curves comparing IFACE with MaSIF (mean (M) and mean–max (X) pooling), SurfaceID (mean (M) and mean–max (X) pooling), Laplace–Beltrami (LB), and JSD distances for discriminating intra-family from inter-family protein pairs. Performance is summarized by Average Precision (AP) and ROC–AUC. IFACE achieves the best overall classification performance among all evaluated methods. (c) Leave-one-feature-out ablation comparing correspondence-based IFACE with matched JSD distances. Removing the structural descriptor produces the largest overall decrease in IFACE performance, highlighting the dominant contribution of intrinsic geometry to family-level discrimination. Among the chemical descriptors, removing hydrogen-bond propensity causes the largest performance decrease, whereas removing electrostatics yields a marginally higher AP than the full feature set.

Beyond the distribution-based (JSD) and intrinsic spectral Laplace–Beltrami baselines, we additionally considered two state-of-the-art learning-based surface representation methods for the family-level benchmark. We used the released pretrained models of MaSIF-search^69^ and SurfaceID^46^ without additional training or fine-tuning to extract local descriptors from the protein surfaces. Since both methods were designed to learn local surface representations rather than whole-protein descriptors, their descriptors were aggregated to enable global protein comparison.

For the learning-based baselines, a fixed-length representation was constructed for each protein surface by aggregating the local patch descriptors produced by the pretrained models. MaSIF-search, trained for protein–protein interface prediction, produces an 80-dimensional descriptor for each surface patch, whereas SurfaceID, trained by self-supervised learning, produces a 512-dimensional descriptor. Because the number of patches varies among proteins, we evaluated two aggregation schemes. In the Mean variant, patch descriptors were averaged over the surface. In the Mean-Max variant, the element-wise mean and maximum were concatenated, yielding 160-dimensional MaSIF-search and 1024-dimensional SurfaceID representations. Pairwise protein distances were then defined as Euclidean distances between these global descriptors.

Figure 5 summarizes the binary classification performance and feature ablation analysis on the family benchmark. The precision–recall (Fig. 5a) and ROC (Fig. 5b) curves demonstrate that IFACE achieves the highest overall performance (AP = 0.95, ROC-AUC = 0.98), outperforming both correspondence-independent baselines, JSD and Laplace–Beltrami (LB), and the learning-based surface descriptors MaSIF Mean (MaSIFM), MaSIF Mean-Max (MaSIFX), SurfaceID Mean (SurfIDM), and SurfaceID Mean-Max (SurfIDX).

Although these learned descriptors were originally developed for local patch comparison and descriptor retrieval rather than whole-protein family classification, they provide useful baselines for assessing how well pretrained local representations transfer to global protein comparison. Unlike the conformer benchmark, where global feature distributions alone are sufficient to distinguish different protein identities, family-level discrimination benefits from preserving the spatial organization of geometric and physicochemical features across complete surfaces.

The feature contribution analysis (Fig. 5c) further supports this observation. In contrast to the conformer benchmark, removing the electrostatic descriptor substantially improves JSD performance, indicating that the marginal electrostatic distribution is comparatively poorly conserved within these protein families and may obscure family-level similarity. Removing electrostatics from IFACE produces only a negligible improvement relative to the full distance. Furthermore, removing either the structural or mean-curvature descriptor causes a larger decrease in IFACE performance than removing any individual chemical descriptor, indicating that geometry plays a more prominent role in family-level discrimination than in conformer recognition. Interestingly, for IFACE, the geometry-only representation (AP = 0.731, ROC-AUC = 0.951) performs noticeably better than the chemical-only representation (AP = 0.628, ROC-AUC = 0.840).

In contrast, combining these complementary sources of information through the full correspondence-aware IFACE representation increases the AP compared with using either modality alone. Although the full IFACE performance is only negligibly lower than that obtained without the electrostatic descriptor, the full representation remains among the best-performing feature configurations, demonstrating that geometric and chemical information provide complementary contributions to family-level discrimination.

A similar trend is observed for JSD, although both its geometry-only and chemical-only representations perform substantially worse than their IFACE counterparts. These observations indicate that explicit surface correspondences preserve complementary geometric and physicochemical information that is largely lost when descriptors are reduced to global feature distributions. Overall, the family benchmark demonstrates that correspondence-aware integration of geometry and chemistry becomes increasingly important as the comparison task shifts from distinguishing different protein identities to recognizing relationships among homologous proteins across diverse families.

We therefore ask whether the protein families emerge as coherent clusters in the full distance space. Table II summarizes both the peak clustering performance and the robustness of each distance measure across a diverse range of clustering strategies. To avoid bias toward a particular clustering algorithm, we evaluated four hierarchical linkage methods (single, complete, average, and weighted) together with a broad collection of Classical Multidimensional Scaling (CMDS) pipelines. The CMDS evaluation included both fixed-dimensional embeddings (4, 5, 6, 8, 10, 15, and 20 dimensions) and variance-preserving embeddings retaining 80%, 90%, 95%, and 99% of the explained variance, each followed by K-means and Ward clustering. To quantify algorithmic robustness, we report the coefficient of variation (CV = SD/Mean) computed separately for the hierarchical methods, the CMDS-based pipelines, and all clustering settings combined, where a lower CV indicates lower sensitivity to the choice of clustering algorithm.

**TABLE II.**
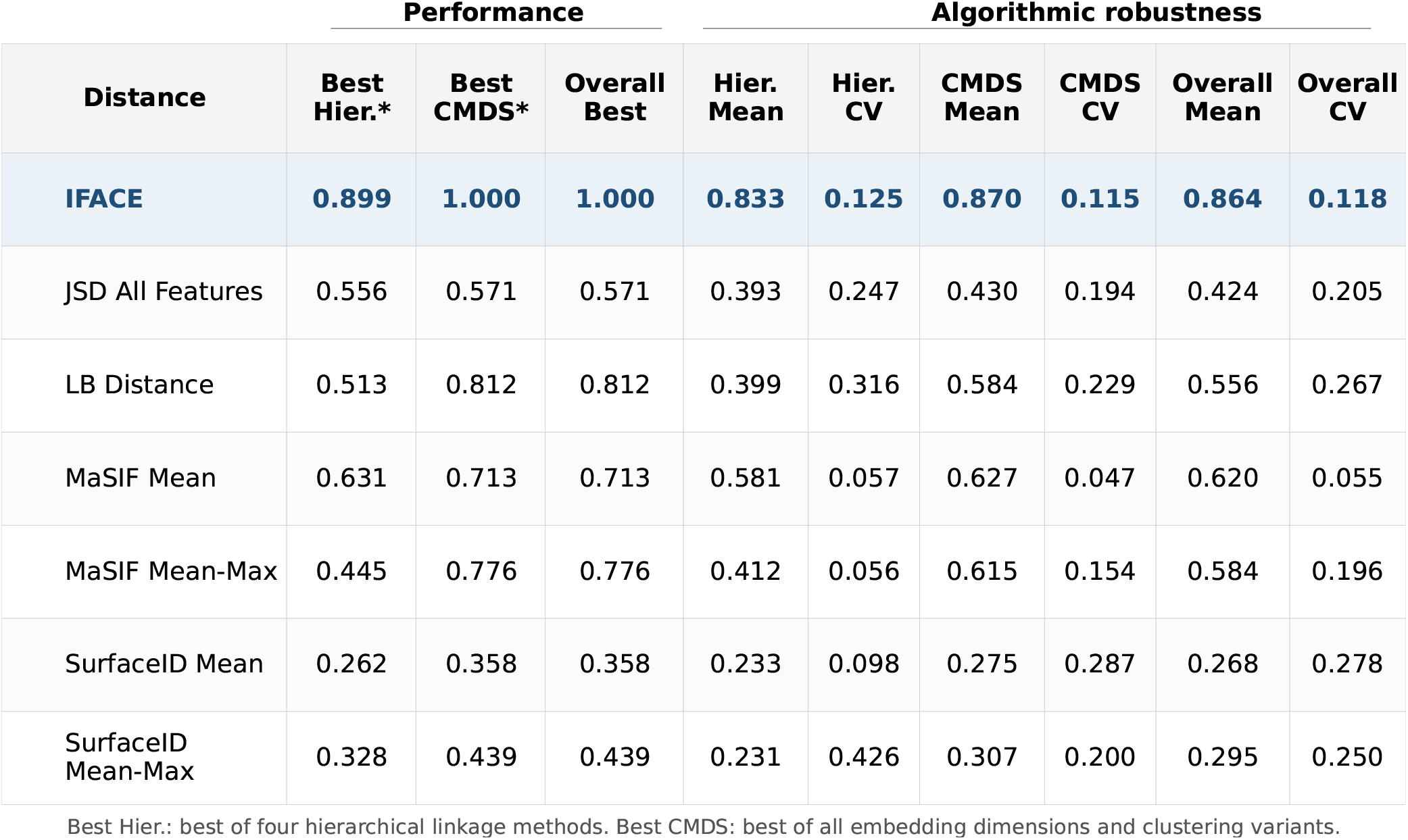
Summary of family-level clustering performance and robustness. Best Hier. (hierarchical) and Best CMDS report the highest Adjusted Rand Index (ARI) obtained using hierarchical and CMDS-based clustering, respectively. Mean and coefficient of variation (CV = SD/Mean) are reported separately for robustness for hierarchical clustering, CMDS-based clustering, and all clustering settings. IFACE outperforms all other methods.

As shown in Table II, IFACE consistently achieves the strongest clustering performance among all evaluated methods. It attains the highest ARI under both hierarchical and CMDS-based clustering strategies, while also delivering the best overall mean ARI. In contrast, the distribution-based JSD distance exhibits substantially weaker clustering performance. Although the Laplace–Beltrami distance outperforms JSD, it remains consistently inferior to IFACE.

Among the learning-based baselines, MaSIF Mean achieves the strongest clustering performance, followed by MaSIF Mean–Max, whereas both SurfaceID variants perform considerably worse. Nevertheless, all learning-based baselines remain well below IFACE. IFACE also demonstrates strong algorithmic robustness, maintaining consistently high clustering accuracy across different clustering algorithms with only limited sensitivity to algorithm selection. Although MaSIF Mean exhibits the lowest variability across clustering strategies, this stability is accompanied by consistently moderate clustering accuracy. Overall, these results show that IFACE provides the best balance of clustering accuracy and algorithmic robustness across a diverse range of hierarchical and CMDS-based clustering approaches.

Consistent with the results of the intra- versus inter-family classification task, the leave-one-feature-out ablation (Table III) demonstrates that the contributions of the individual descriptors are not uniform. Removing the structural descriptor produces the largest degradation in clustering performance, reducing the overall mean ARI from 0.864 to 0.705, indicating that intrinsic surface geometry provides the largest individual contribution among the descriptors.

**TABLE III.**
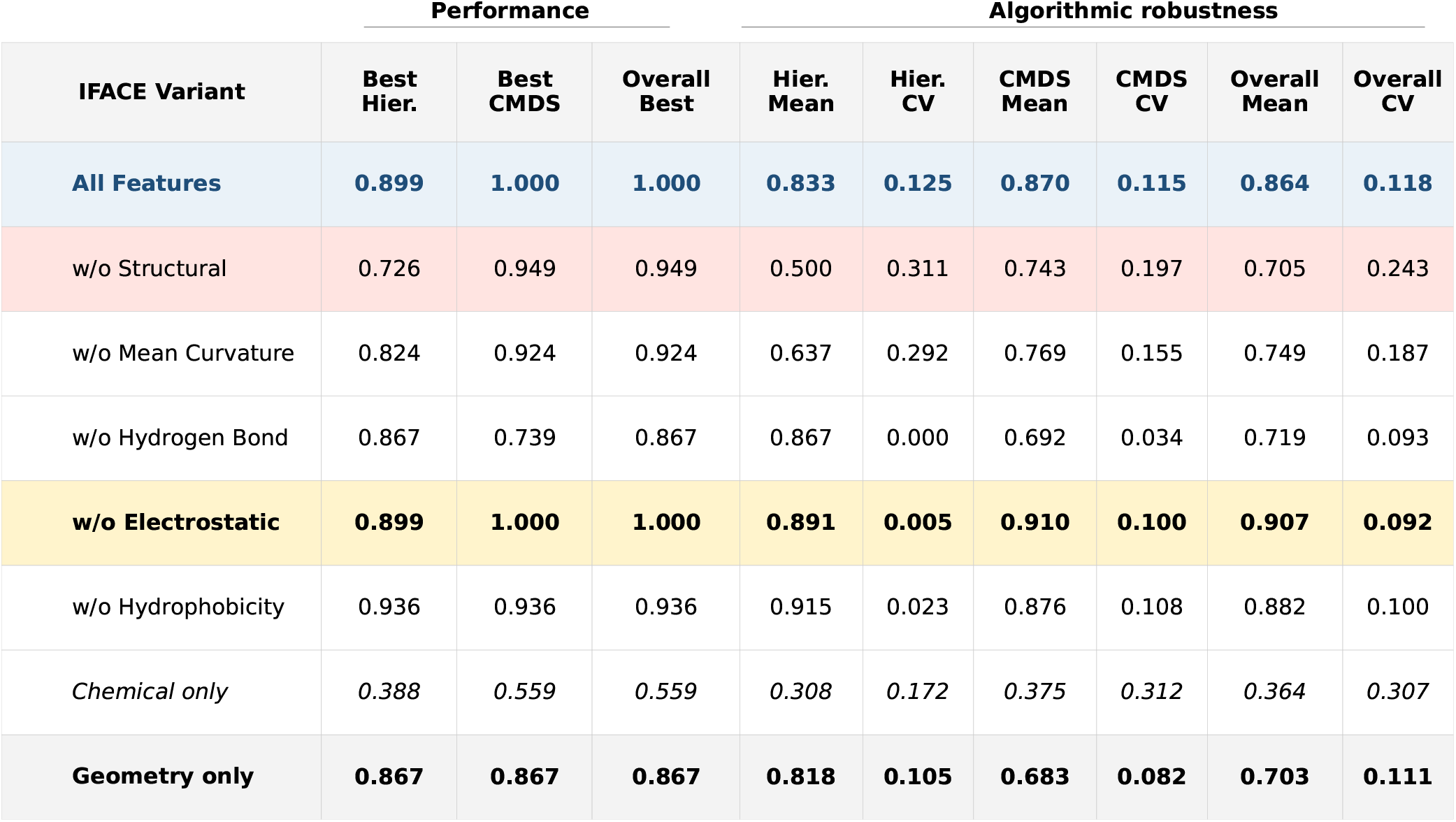
Summary of clustering performance and robustness of IFACE distance ablation on the family benchmark. Best Hier. (hierarchical) and Best CMDS report the highest Adjusted Rand Index (ARI) obtained using hierarchical and CMDS-based clustering, respectively. Mean and coefficient of variation (CV = SD/Mean) summarize performance across hierarchical methods, CMDS-based clustering, and all clustering settings. Removing the electrostatic descriptor modestly improves mean clustering performance, whereas removing the structural descriptor produces the largest degradation.

| IFACE Variant | Performance |  |  | Algorithmic robustness |  |  |  |  |  |
| --- | --- | --- | --- | --- | --- | --- | --- | --- | --- |
|  | Best Hier. | Best CMDS | Overall Best | Hier. Mean | Hier. CV | CMDS Mean | CMDS CV | Overall Mean | Overall CV |
| **All Features** | **0.899** | **1.000** | **1.000** | **0.833** | **0.125** | **0.870** | **0.115** | **0.864** | **0.118** |
| w/o Structural | 0.726 | 0.949 | 0.949 | 0.500 | 0.311 | 0.743 | 0.197 | 0.705 | 0.243 |
| w/o Mean Curvature | 0.824 | 0.924 | 0.924 | 0.637 | 0.292 | 0.769 | 0.155 | 0.749 | 0.187 |
| w/o Hydrogen Bond | 0.867 | 0.739 | 0.867 | 0.867 | 0.000 | 0.692 | 0.034 | 0.719 | 0.093 |
| **w/o Electrostatic** | **0.899** | **1.000** | **1.000** | **0.891** | **0.005** | **0.910** | **0.100** | **0.907** | **0.092** |
| w/o Hydrophobicity | 0.936 | 0.936 | 0.936 | 0.915 | 0.023 | 0.876 | 0.108 | 0.882 | 0.100 |
| *Chemical only* | *0.388* | *0.559* | *0.559* | *0.308* | *0.172* | *0.375* | *0.312* | *0.364* | *0.307* |
| **Geometry only** | **0.867** | **0.867** | **0.867** | **0.818** | **0.105** | **0.683** | **0.082** | **0.703** | **0.111** |

Nevertheless, geometry alone, comprising both intrinsic structural and extrinsic mean-curvature information, does not fully capture protein family similarity. Although the geometry-only variant substantially outperforms the chemical-only variant (overall mean ARI of 0.703 versus 0.364), it remains inferior to the complete IFACE distance (overall mean ARI of 0.864), demonstrating that geometric and physicochemical descriptors provide complementary information for protein family discrimination.

Conversely, the chemical-only variant exhibits consistently weak clustering performance, indicating that physicochemical information alone is insufficient for reliable family-level discrimination within this benchmark. However, removing the electrostatic descriptor modestly increases the overall mean ARI, whereas removing hydrophobicity produces a smaller improvement.

Overall, the complete IFACE distance performs strongly on both the conformer and family benchmarks, indicating that integrating intrinsic geometry, extrinsic geometry, and multiple physicochemical descriptors yields a robust protein-surface comparison distance without task-specific feature selection. These conclusions are further supported by the cluster-purity results (Supplementary Tables), where IFACE generally achieves the highest or among the highest purity scores across the evaluated clustering algorithms.

Furthermore, the corresponding leave-one-feature-out ablation for the distribution-based JSD distance (see Supplementary Information) is generally weaker than its IFACE counterpart, indicating that global feature distributions do not reproduce the same family-level organization. Complete leave-one-feature-out and feature-combination ablation studies are provided in the Supplementary Information.

Taken together, these results show that combining geometry with physicochemical surface fields enables IFACE to produce a stable and discriminative family-level organization within this benchmark.

#### EFFECT OF SURFACE RESOLUTION

To assess whether the surface discretization was sufficient, we repeated the family- and conformer-level classification and clustering analyses using surfaces downsampled to 1,000 vertices and compared the results with those obtained using 3,000 vertices. Performance remained comparable across the two resolutions, indicating that the lower-resolution meshes preserve the geometric and physicochemical information required to distinguish protein families and conformational states.

At 1,000 vertices, IFACE maintained performance comparable to that obtained at 3,000 vertices. The conformer benchmark yielded an AP of 0.91 and a ROC-AUC of 0.92, while the family benchmark yielded an AP of 0.97 and a ROC-AUC of 0.99. Hierarchical clustering achieved an ARI of 0.90, and the CMDS-based clustering results were similarly strong across several embedding dimensions. Thus, 1,000-vertex surfaces provide an adequate representation for these analyses while substantially reducing the computational cost of the correspondence calculation (see Section Scaling)

#### P450 HEME-POCKET-ACTIVE SITE TRANSFER EVALUATION

To directly evaluate whether the computed point-to-point correspondences correctly transfer functional pocket regions between homologous proteins, we define a vertexwise pocket-transfer score.

For each protein, we first identify the surface vertex nearest to the bound heme cofactor and use it as the center of a heme-centered pocket region. The strict pocket is defined as the set of surface vertices whose graph-geodesic distance from this vertex is at most 20 Å. This region is highly localized, covering only a small fraction of the total protein surface. For the relaxed evaluation, the target acceptance region is expanded to a graph-geodesic radius of 22 Å, allowing small localization errors near the pocket boundary while preserving the same overall functional neighborhood.

For each directed protein pair, every source-pocket vertex is mapped to its ranked IFACE correspondences on the target surface. Under the Top-1 evaluation, only the highest-scoring correspondence is considered. Under the Top-2 evaluation, which is the primary setting used in the main analysis, a source-pocket vertex is counted as successfully transferred if either of its two highest-scoring target correspondences lies within the target pocket. Each source-pocket vertex contributes at most one successful transfer, irrespective of how many candidate correspondences fall inside the target pocket.

The pocket transfer score is defined as the fraction of source-pocket vertices that are successfully localized within the target pocket. Because the score is normalized by the number of source-pocket vertices, it measures how completely the source functional region is recovered on the target surface and permits comparison across source pockets of different sizes. The strict and relaxed target definitions are reported separately because expanding the target region changes the acceptance criterion.

To summarize transfer performance across protein pairs, we additionally define a pocket transfer hit. A directed transfer is considered successful when at least 50% of the source-pocket vertices are correctly localized within the target pocket. This majority-recovery criterion serves as the primary operating threshold in the main analysis.

Recovering at least half of the source pocket provides an operational indication that the corresponding heme-centered region has been substantially localized on the target surface, although the 50% threshold should not be interpreted as a biologically validated cutoff. In addition, to assess sensitivity to the chosen stringency, we report success rates over recovery thresholds ranging from 30% to 90%.

Figure 6 summarizes pocket-transfer performance across the eight P450 proteins by varying the minimum pocket transfer score required for a successful mapping. At the primary 50% threshold, 47 of the 56 directed mappings (83.9%) are successful under Strict Top-1, increasing to 50 mappings (89.3%) under Strict Top-2. Under the relaxed evaluation, the corresponding success rates are 53 of 56 mappings (94.6%) for Top-1 and 54 of 56 mappings (96.4%) for Top-2.

**FIG. 6.**
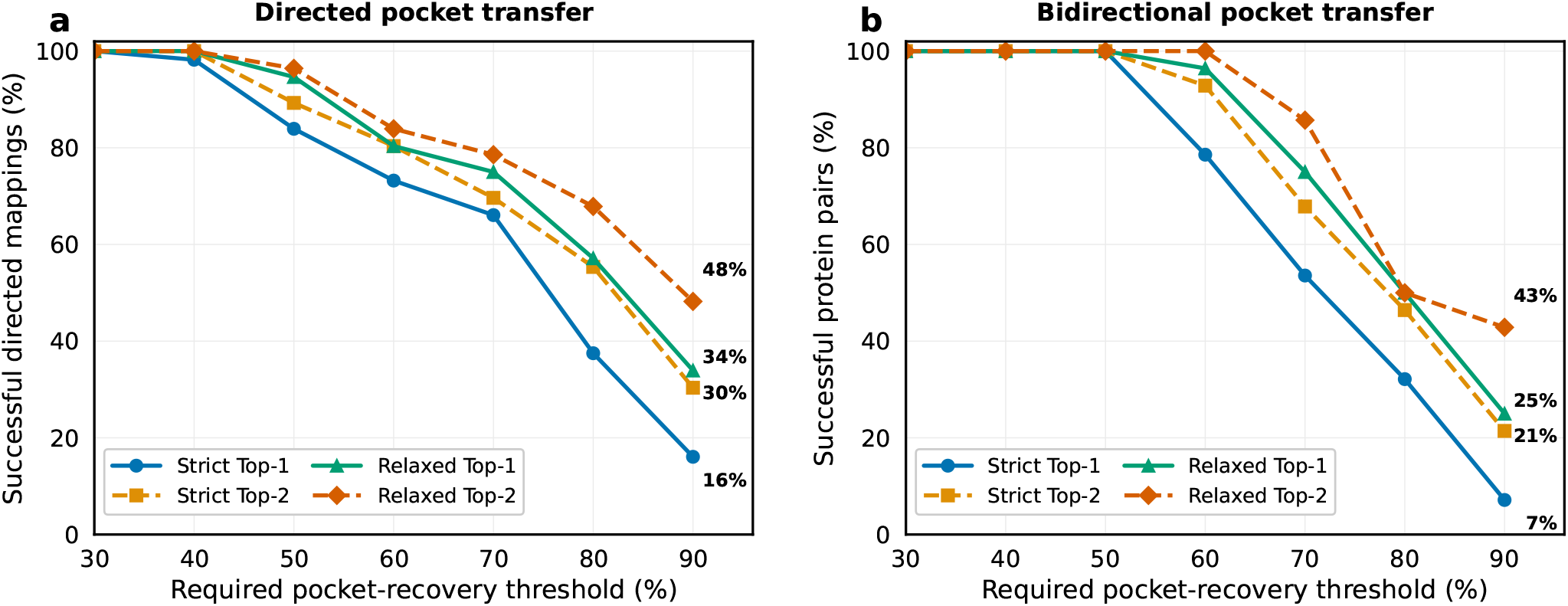
Threshold-dependent pocket-transfer performance for the heme-centered benchmark. The horizontal axis denotes the minimum required pocket-recovery threshold, while the vertical axis reports the percentage of successful mappings satisfying that threshold. (a) Percentage of successful directed transfers among all 56 ordered protein mappings. (b) Percentage of successful bidirectional protein pairs among the 28 unordered pairs, where a pair is counted only if the mean recovery over both mapping directions exceeds the specified threshold. Overall, relaxed-pocket evaluation consistently outperforms the strict-pocket criterion, and allowing Top-2 correspondences further improves transfer success relative to Top-1 matching.

For the 28 unordered protein pairs, all 28 pairs (100%) satisfy the 50% threshold when recovery is averaged over both mapping directions, under either the strict or relaxed pocket definition and with either Top-1 or Top-2 correspondences. As expected, the success rates decrease monotonically as the required pocket recovery becomes more stringent. The differences between the evaluation settings become more pronounced above the 60% threshold.

At the most stringent 90% threshold, Strict Top-1 and Top-2 succeed for 9 and 17 of the 56 directed mappings, corresponding to 16.1% and 30.4%, respectively. Relaxed Top-1 and Top-2 succeed for 19 and 27 mappings, corresponding to 33.9% and 48.2%, respectively. Among the 28 unordered pairs, Strict Top-1 and Top-2 succeed for 2 and 6 pairs (7.1% and 21.4%), whereas Relaxed Top-1 and Top-2 succeed for 7 and 12 pairs (25.0% and 42.9%). Across the evaluated thresholds, Top-2 and relaxed-pocket evaluations yield higher success rates than Top-1 and strict-pocket evaluations, respectively, demonstrating the increased recovery obtained by considering an additional high-ranking correspondence and a slightly expanded target region.

These results should be interpreted in the context of heme-centered P450 active sites, which are typically buried or semi-buried cavities connected to solvent or membrane through access channels rather than simple exposed surface patches. In such systems, accurate transfer requires not only locating the correct heme-proximal region but also reconstructing the geometry and extent of the surrounding cavity and access channels. Recovering at least 50% of the source-pocket vertices is therefore a useful indicator that the correct functional region has been localized, whereas higher thresholds demand increasingly faithful reconstruction of the pocket extent.

The higher performance obtained with the relaxed pocket definition is consistent with many errors occurring near pocket boundaries or small cavity offsets, but does not by itself prove that all errors are boundary-localized. Similarly, the advantage of Top-2 over Top-1 indicates that allowing an additional candidate correspondence improves recall under this evaluation, although the threshold plots alone do not establish the precise structural reason why the second-ranked candidate helps.

#### CORRESPONDENCE-QUALITY ANALYSIS

To assess the quality of the recovered correspondences, we evaluate neighborhood preservation using geodesic neighborhoods (Fig. 7). For each source vertex, a neighborhood is defined by a geodesic ball of radius 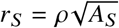, where *A*_*S*_ is the total surface area of the source mesh. The corresponding target neighborhood is centered at the target vertex to which the source vertex is mapped and has radius *R* = *αr*_*T*_, where 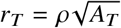 and *A*_*T*_ is the total surface area of the target mesh.

**FIG. 7.**
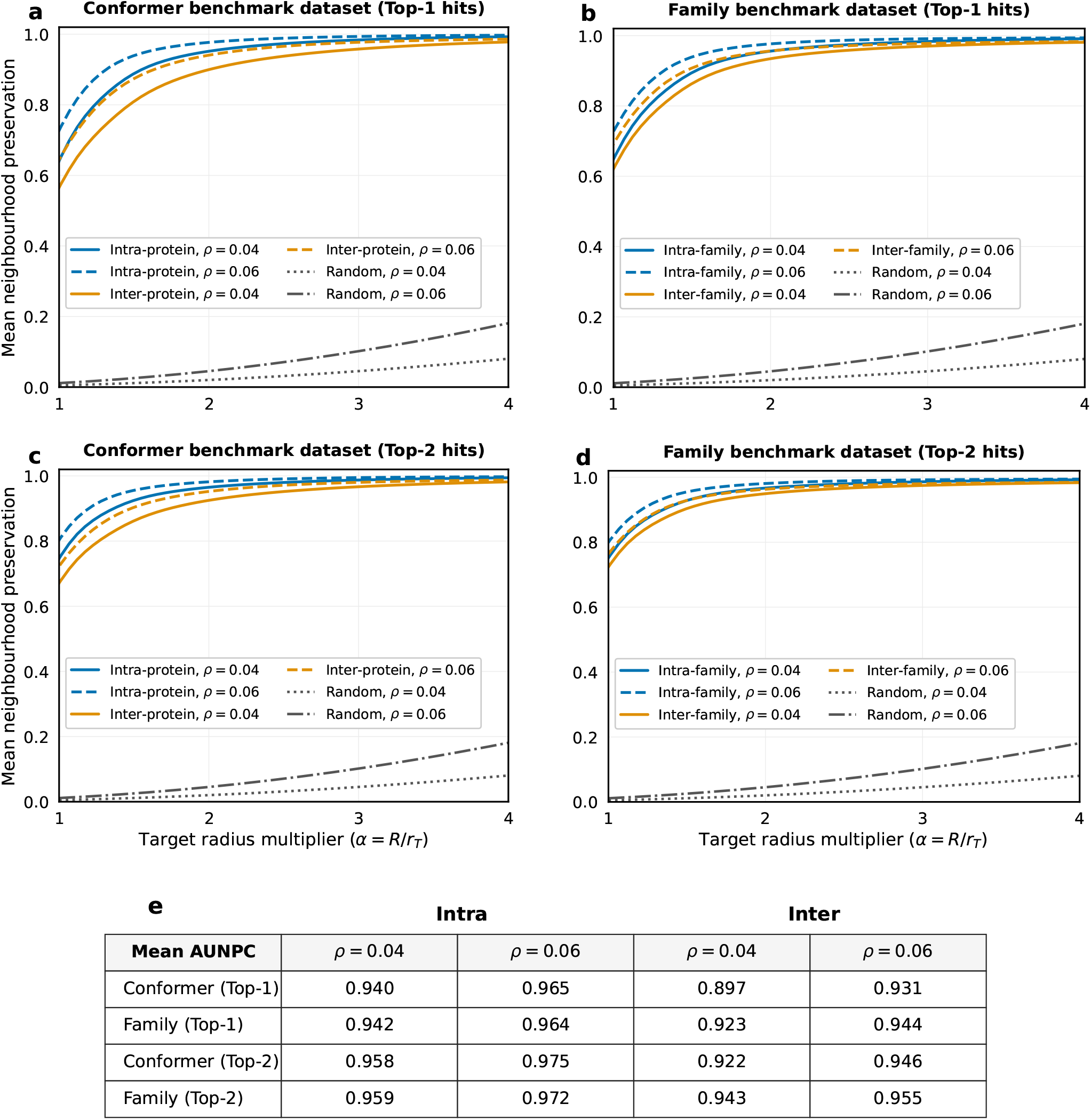
Neighborhood preservation analysis of the recovered protein surface correspondences. **(a)** Conformer benchmark using hard correspondences. **(b)** Family benchmark using hard correspondences. **(c)** Conformer benchmark using top-2 relaxed correspondences. Family benchmark using top-2 relaxed correspondences. **(e)** Mean Area Under the neighborhood Preservation Curve (AUNPC) for the four experimental settings. Panels (a)–(d) show the mean neighborhood preservation as a function of the target radius multiplier *α* for *ρ* = 0.04 and *ρ* = 0.06.

The neighborhood preservation score is defined as the fraction of mapped source neighbors that lie within the target geodesic ball, averaged over all source vertices. We report this quantity as a function of the target radius multiplier *α* for *ρ* = 0.04 and *ρ* = 0.06, corresponding to neighborhoods containing approximately 12 and 30 vertices on average.

At *α* = 1, the source and target neighborhoods are defined at the same area-normalized geodesic scale. Since the source and target meshes have comparable sampling densities and similar numbers of vertices, they contain approximately the same number of vertices on average. Thus, *α* = 1 provides the natural reference point for neighborhood preservation, while *α* > 1 measures the additional target radius required to recover the mapped neighborhood.

To summarize the entire curve with a single scalar, we compute the Area Under the neighborhood Preservation Curve (AUNPC) by integrating the neighborhood preservation curve over *α* ∈ [1, 4] and normalizing by the integration interval. We report results using both hard correspondences, obtained by assigning each source vertex to its highest-probability target vertex, and a top-2 relaxed evaluation, in which a neighboring source vertex is considered preserved if either of its two highest-probability target candidates lies within the target geodesic ball.

Figure 7 demonstrates that IFACE recovers correspondences that preserve local surface neighborhoods across both the conformer and family benchmarks. In all cases, neighborhood preservation increases monotonically with the target radius multiplier *α* and approaches unity as the target neighborhood expands, indicating that mapped neighbors remain spatially coherent under the recovered correspondences.

As expected, intra-protein and intra-family pairs consistently exhibit higher neighborhood preservation than interprotein and inter-family pairs, respectively, particularly around the natural operating point *α* = 1, where the source and target neighborhoods are defined at the same normalized geodesic scale. This separation indicates that IFACE produces more locally consistent correspondences for same-protein conformers and proteins from the same family than for unrelated proteins, while the gradual convergence of the curves at larger *α* reflects the expected recovery of neighborhoods as the target radius is expanded.

The top-2 relaxed evaluation consistently improves neighborhood preservation over the hard assignment on both benchmarks, indicating that the soft correspondence matrix retains additional geometrically consistent candidate matches beyond the single best assignment. Nevertheless, the improvement is modest, suggesting that the highest-probability correspondence already captures most of the local neighborhood structure recovered by IFACE.

These observations are summarized quantitatively in Fig. 7(e), which reports the mean Area Under the neighborhood Preservation Curve (AUNPC). Across all evaluated settings, AUNPC values exceed 0.89 and consistently increase under the top-2 relaxed evaluation.

For example, on the family benchmark, the mean AUNPC increases from 0.942 to 0.959 for *ρ* = 0.04 and from 0.964 to 0.972 for *ρ* = 0.06 for intra-family pairs, while the corresponding inter-family values increase from 0.923 to 0.943 and from 0.944 to 0.955, respectively.

Similarly, on the conformer benchmark, the mean AUNPC increases from 0.940 to 0.958 (*ρ* = 0.04) and from 0.965 to 0.975 (*ρ* = 0.06) for intra-protein pairs, while the corresponding inter-protein values increase from 0.897 to 0.922 and from 0.931 to 0.946, respectively.

Overall, these consistently high AUNPC values indicate that IFACE recovers correspondences with strong local geometric consistency, while the gains achieved by the top-2 evaluation indicate that the soft correspondence matrix retains additional plausible local matches beyond the highest-probability assignment.

#### SCALING

To evaluate the computational scaling of IFACE, we measured the runtime and peak memory consumption of the complete pipeline for a representative protein pair while varying the mesh resolution. Both the source and target surfaces were uniformly downsampled to approximately 1,000, 2,000, 3,000, and 4,000 vertices, while all algorithmic parameters and optimization settings were held fixed. Figures 8(a) and 8(b) show the total runtime and peak resident memory usage, respectively, as functions of the number of vertices per surface. The dashed curves denote leastsquares power-law fits of the form (*y* = *CN*^*p*^). Over the range of mesh resolutions considered, the fitted relations were (*T* (*N*) ≈ 3.17 × 10^−7^ *N*^2.67^) for runtime and (*M* (*N*) ≈ 1.24 × 10^−4^ *N*^1.21^) for peak memory usage, where runtime is measured in seconds and memory in gigabytes.

**FIG. 8.**
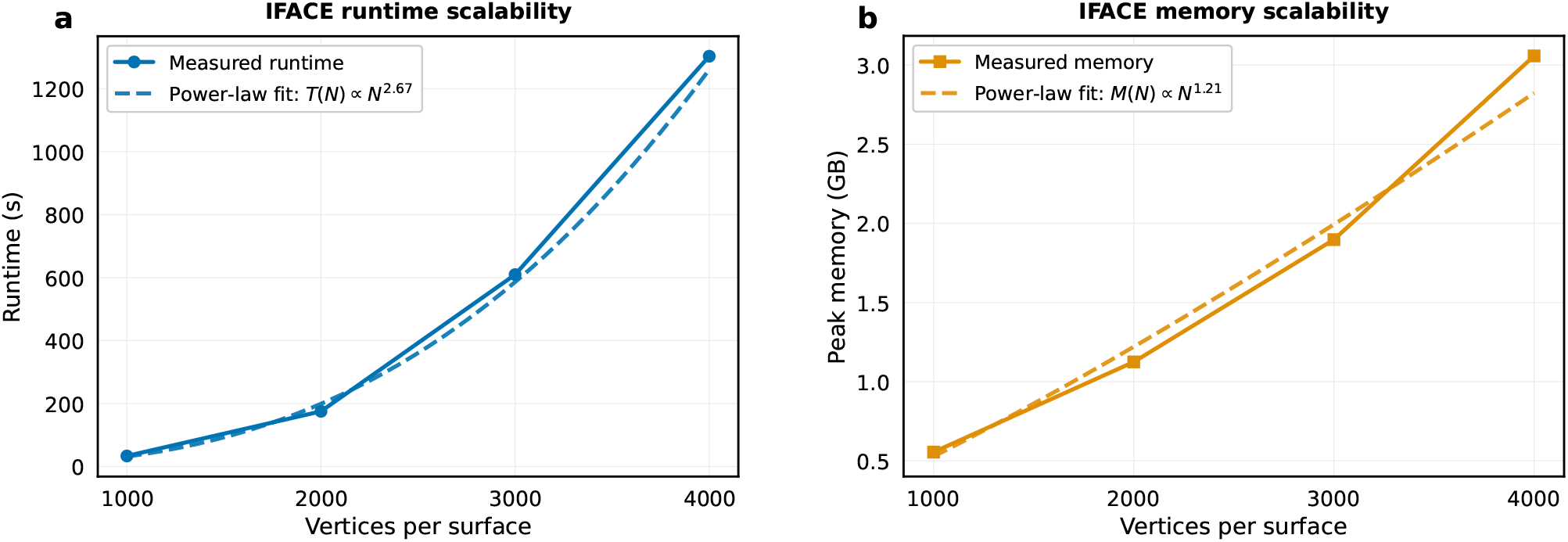
Scalability analysis of IFACE with increasing mesh resolution. A representative protein pair was uniformly downsampled to different mesh resolutions while keeping all algorithmic parameters fixed. **(a)** Total runtime of the complete IFACE pipeline, including coupling optimization and distance calculation. **(b)** Peak resident memory usage during execution. Solid curves show the measured values, while dashed curves represent least-squares power-law fits of the form *y* = *CN*^*p*^, illustrating the empirical scaling of IFACE with mesh resolution (number of vertices per surface).

## DISCUSSION

Protein surfaces are naturally represented as coupled geometric–chemical objects, and their similarity is most directly defined through correspondence between surfaces. IFACE implements this principle by constructing an optimal-transport coupling, yielding both an explicit soft correspondence and a symmetric geometric–chemical distance. Similarity is therefore determined by how geometry and physicochemical fields align across complete surfaces, rather than by comparing them separately.

Across the proteins examined here, surface-based distances distinguished intra-protein conformational variability from inter-protein divergence more effectively than global fold-based similarity measures. Incorporating the coupled organization of surface geometry and physicochemical fields reduced the overlap between the intra-protein and inter-protein distance distributions. The stronger performance of JSD in this binary identity test shows that aggregate surface-feature distributions can be sufficient when the task is simply to distinguish one protein from another. This indicates that protein identity is reflected not only in the global fold but also in the spatial arrangement of geometric and physicochemical properties across the molecular surface.

At the family level, IFACE coherently organized proteins from six families and separated the outlier. Geometric distances captured shared surface shape and curvature organization, whereas chemical distances captured complementary physicochemical similarity. Their combination produced stable separation between intra-family and interfamily protein pairs. The contribution of mean curvature further indicates that extrinsic surface organization contains information not fully represented by intrinsic geometry or chemical fields alone.

Compared with the baseline methods, IFACE provided stronger family-level organization and separation of intrafamily and inter-family pairs. The weaker performance of JSD-based distances indicates that similarity between marginal feature distributions is insufficient when the spatial arrangement of surface properties is discarded. Likewise, the Laplace–Beltrami spectral distance captures global intrinsic geometry but does not incorporate physicochemical organization. The learning-based MaSIF and SurfaceID descriptors also showed weaker family-level discrimination on the present dataset. This may reflect their design for task-specific local surface comparisons rather than for defining a general distance between complete protein surfaces. These comparisons support the importance of jointly evaluating geometry and physicochemical fields through explicit correspondence.

The results indicate that family-level relationships are reflected in coupled surface organization. Proteins belonging to the same family retained related geometric and physicochemical surface patterns despite sequence variation and differences in species of origin. The present analysis does not establish a direct mechanistic relationship between the IFACE distance and biological function. Nevertheless, the consistent family-level separation suggests that coupled surface organization may provide a useful indicator of functional relatedness.

An important advantage of IFACE is that the correspondence itself is an output of the comparison. The resulting mapping supports both global analyses, including classification and clustering, and local analyses of corresponding pockets and surface regions. The neighborhood-preservation and pocket-transfer analyses further demonstrate that the correspondence preserves coherent local surface organization. IFACE therefore provides both a global similarity measure and an interpretable mapping without requiring task-specific supervision.

Several limitations warrant consideration. The present implementation uses a limited set of scalar surface fields, and the benchmarks contain a limited number of protein families and conformational ensembles. Broader validation across more diverse proteins and functional surface regions will be required to establish generality. Performance remained comparable when the surface resolution was reduced from 3,000 to 1,000 vertices, indicating that the principal geometric–chemical organization is retained at substantially lower computational cost.

Nevertheless, the computational cost increases with surface resolution, motivating future development of sparse, multiscale, or accelerated transport formulations. Extensions to vector- or tensor-valued fields and alternative surface representations may further refine the correspondence and distance definitions. Future studies can test whether the same framework supports large-scale retrieval and comparison of binding pockets, interaction interfaces, and other functionally annotated surface regions.

Overall, IFACE formulates protein-surface similarity as a coupled geometric–chemical comparison mediated by explicit correspondence between surfaces. By providing both a global distance and an interpretable local mapping, it offers a physically explicit, mathematically principled, and unsupervised framework for comparing functional protein surfaces.

## METHODS

In this section, we describe how distances between protein surfaces are constructed within the IFACE framework. The key step is the identification of a soft correspondence between two surfaces, encoded in an optimal coupling matrix that relates their surface vertices.

### Optimal Coupling Matrix

We consider two protein surfaces, *S*_*α*_ and *S*_*β*_, represented as triangulated meshes and endowed with *m* surface feature fields, denoted 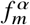 and 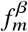. Our central objective is to relate these two surfaces in a manner that respects *both* their geometric structure and surface-field organization. To achieve this, we introduce a probabilistic, or *soft*, correspondence between the surfaces, encoded by a *coupling matrix* **P**. Each matrix element *P*_*i j*_ quantifies the degree to which vertex *i* on *S*_*α*_ corresponds to vertex *j* on *S*_*β*_. By avoiding a rigid one-to-one assignment, this formulation permits a flexible comparison of surface regions and naturally accommodates structural variability and chemical heterogeneity.

The coupling matrix is constrained by prescribed marginal distributions over the vertices of each surface, *ρ*^*α*^ and *ρ*^*β*^, such that 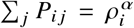 and 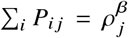. These marginals assign weights to surface elements based on local surface area and the values of the surface feature fields. They therefore act as nonuniform vertex weights in the correspondence constraints. The explicit construction of these distributions is described in the Supplementary Information.

The optimal coupling matrix **P** is determined by balancing two complementary contributions. The first is a *field term*, which compares the surface feature fields defined on the two surfaces. The second is a *structural term*, which enforces geometric compatibility between surface shapes. The optimal correspondence emerges from reconciling these two contributions under the marginal constraints above, yielding a coupling that reflects both field similarity and geometric coherence. We now define these two terms explicitly.

### Field Term

The field term ℱ quantifies the mismatch between feature fields on the two surfaces. For optimization purposes, it is defined as

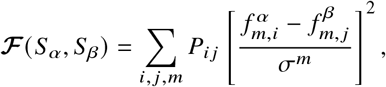

where *σ*^*m*^ denotes the standard deviation of the *m*^th^ feature field, computed using values from both surfaces. This normalization places all fields on a comparable scale and promotes the matching of vertices with similar surface-field values.

### Structural Term

The structural term enforces consistency between the intrinsic geometries of the two surfaces. To this end, we construct for each surface *S* its geodesic distance matrix 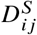 on the unit-normalized mesh, supplemented by a smoothing kernel 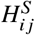 that incorporates global geometric information. These are combined as 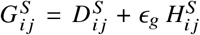 with *S* ∈ {*S*_*α*_, *S*_*β*_}. Here, the smoothing kernel is 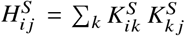, with 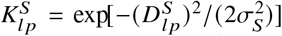. Thus, *H*^*S*^ encodes smoothed, long-range geometric correlations derived from the geodesic matrix.

Using these definitions, the structural discrepancy between surfaces *S*_*α*_ and *S*_*β*_ is given by

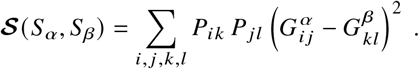

In our analysis, *σ*_*S*_ is chosen as a fixed fraction of the maximum geodesic distance, 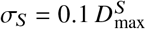, and the weight *ϵ*_*g*_ is set to 0.1 to ensure that the global contribution remains a perturbative correction of the geodesic structure. In the limit *ϵ*_*g*_ = 0, this term reduces to the classical Gromov– Wasserstein objective, with 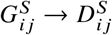.^70,71^

### Optimization

To determine the coupling matrix, we minimize the following entropic-regularized objective function:

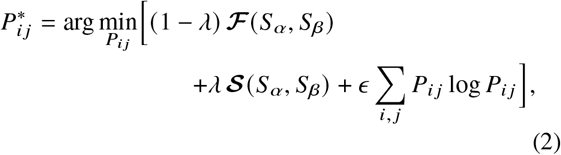

where the minimization is subject to the constraints 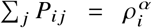 and 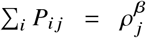. The entropic term −*ϵ* ∑ _*i, j*_ *P*_*i j*_ log *P*_*i j*_ plays a distinct and complementary role to the geometric regularization controlled by *ϵ*_*g*_. While *ϵ*_*g*_ smooths the *metric structure* entering the structural term, the entropy promotes diffuseness of the coupling itself, suppressing spurious sharp correspondences and stabilizing the optimization landscape. In practice, this regularization improves convergence and prevents the solution from becoming trapped in shallow local minima.

To find the optimal coupling, we proceed in several stages. We first obtain an initial alignment of the meshes using a combination of rigid and feature-field-aware non-rigid alignment, based on mean curvature, hydrophobicity, hydrogen bonding, and electrostatic potential. This alignment is converted into an initial coupling matrix using a Gaussian kernel followed by Sinkhorn normalization^72,73^ to enforce the marginal constraints. The resulting coupling serves as a warm start for minimizing the entropically regularized objective. Finally, the entropically regularized solution is used to initialize a second optimization with *ϵ* = 0, yielding the final coupling matrix 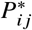. Full algorithmic details are provided in the Supplementary Information.

We next describe how distances between protein surfaces are constructed from the resulting coupling matrix.

### IFACE Distance

We construct distances between protein surfaces by leveraging the optimal coupling to compare both surface feature fields and geometric structure. The guiding principle is to use only the most reliable portions of the correspondence: for each vertex, we retain the two most confidently mapped vertices on the opposing surface. Feature values are transported across the coupling, evaluated at the mapped vertices, and compared using the *ℓ*^1^ norm, *i*.*e*., absolute value. The choice of the *ℓ*^1^ norm is deliberate: it suppresses the influence of outliers that can arise from discretization effects, where feature values may vary unevenly across meshes of differing resolution, thereby yielding a more robust estimate of feature-field dissimilarity.

To ensure symmetry, all comparisons are performed *bidirectionally*. In addition, prior to computing distances, feature fields are mildly regularized by local smoothing. For each vertex *i*, the field value is updated using its neighborhood according to

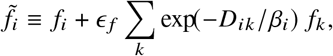

where *D* is the geodesic distance matrix and *β*_*i*_ is chosen as the characteristic distance of the 2-ring neighborhood of vertex *i*. Throughout this work, we fix *ϵ* _*f*_ = 0.1. In the following expressions, 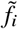 denotes the smoothed field.

Using these smoothed fields, the feature-field distance *D*_field_ is defined as

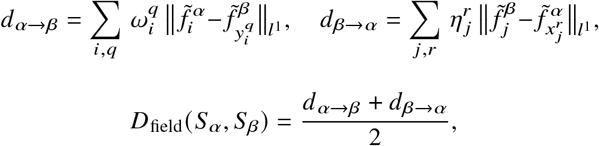

where 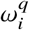 and 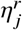 are confidence weights associated with the vertex mappings 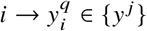 and 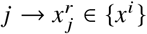, respectively. These weights are normalized such that 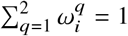 and 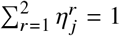, ensuring that only the relative confidence between the two dominant correspondences enters the distance.

The structural distance is constructed analogously using a bidirectional comparison of the structural matrices *G*^*α*^ and *G*^*β*^, which are computed from the original protein meshes rather than the unit-normalized meshes. It is defined as

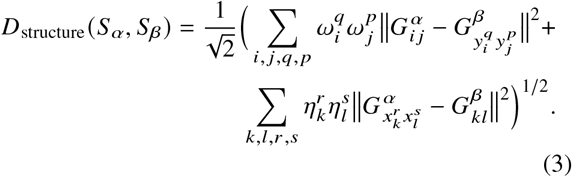

All notation follows the definitions introduced above. This construction yields a symmetric structural distance that reflects discrepancies in intrinsic geometry while remaining tightly coupled to the most confident regions of the surface correspondence.

To combine the geometric and chemical information into a unified surface-comparison distance, we first normalize each distance component before aggregation. The normalization bounds are computed from a reference dataset of 130 proteins spanning both the protein-family and conformer benchmarks. This reference dataset provides a broad distribution of distances, allowing the different components to be placed on comparable numerical scales and preventing any component from dominating solely because of its magnitude.

For each surface scalar field *m*, the range-normalized field distance is defined as

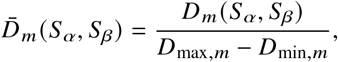

where *D*_*m*_ (*S*_*α*_, *S*_*β*_) is the bidirectional field distance defined above, and *D*_min,*m*_ and *D*_max,*m*_ are the minimum and maximum values, respectively, of that distance component in the reference dataset. The empirical range *D*_max,*m*_ − *D*_min,*m*_ is used as a scale factor; the reference minimum is not subtracted, thereby preserving zero raw distance. The structural distance is normalized independently as

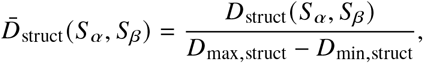

where *D*_min,struct_ and *D*_max,struct_ are the corresponding reference bounds for the structural distance.

Because mean curvature is a geometric scalar field, the geometric distance is defined as the sum of the normalized structural distance and the normalized mean-curvature field distance,

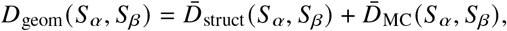

where 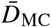 denotes the range-normalized bidirectional distance between the mean-curvature fields.

The chemical distance is defined as the sum of the normalized physicochemical field distances,

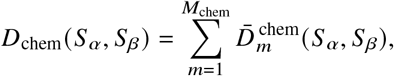

where *M*_chem_ is the number of physicochemical surface fields. In the present formulation, these fields are electrostatic potential, hydrogen-bond propensity, and hydrophobicity.

Finally, the IFACE distance is defined as the sum of the geometric and chemical distances,

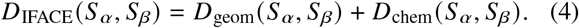

## Data and Software Availability

The dataset used for the analysis can be freely downloaded from Zenodo: https://doi.org/10.5281/zenodo.21760918. A working implementation of the IFACE framework, including surface preprocessing, coupling computation, and distance evaluation, is available on GitHub: https://github.com/ihswami/iface.

## AUTHOR CONTRIBUTIONS

H.S. conceived the differential-geometric approach underlying IFACE and developed the IFACE methodology presented in this work, including its mathematical formulation and computational implementation, wrote the software, performed the analyses and validation, curated the data, prepared the original draft, and contributed extensively to its revision. J.M.M., J.-P.E., and T.T. conceived the broader project of mapping and comparing protein surfaces. J.M.M. developed software for dataset processing and method evaluation and contributed to data curation, supervision, and writing. J.-P.E. contributed to mathematical and theoretical development, interpretation of the results, scientific supervision, and writing. T.T. guided and supervised the scientific development of the project and contributed to interpretation and writing.

## SUPPORTING INFORMATION

General IFACE framework and probabilistic soft mapping; discrete protein-surface construction and coupling optimization; comments on geometry-based correspondence methods; distance-normalization procedure; complete ablation and clustering tables; and P450 pocket-transfer threshold tables (PDF).

## ACKNOWLEDGMENTS

This work was supported by the National Research Foundation of Korea under Grant No. NRF-RS-2025-00573354, the InnoCORE Bio-MAX program, and the U.S. Office of Naval Research under Award No. N00014-26-1-2401. J.-P.E. was partially supported by SwissMAP.

## REFERENCES

[1] J. Janin and F. Rodier, Protein–protein interaction at crystal contacts, Proteins: Structure, Function, and Bioinformatics 23, 580 (1995).

[2] L. H. Hartwell, J. J. Hopfield, S. Leibler, and A. W. Murray, From molecular to modular cell biology, Nature 402, C47 (1999).

[3] D. S. Goodsell and A. J. Olson, Structural symmetry and protein function, Annual review of biophysics and biomolecular structure 29, 105 (2000).

[4] T. Tlusty and A. Libchaber, Life sets off a cascade of machines, Proceedings of the National Academy of Sciences 122, e2418000122 (2025).

[5] E. Weinreb, J. M. McBride, M. Siek, J. Rougemont, R. Renault, Y. Peleg, T. Unger, S. Albeck, Y. Fridmann-Sirkis, S. Lushchekina, et al., Enzymes as viscoelastic catalytic machines, Nature Physics 21, 787 (2025).

[6] J. M. McBride and T. Tlusty, The physical logic of protein machines, Journal of Statistical Mechanics: Theory and Experiment 2024, 024001 (2024).

[7] K. B. Zeldovich and E. I. Shakhnovich, Understanding protein evolution: from protein physics to darwinian selection, Annu. Rev. Phys. Chem. 59, 105 (2008).

[8] D. A. Liberles, S. A. Teichmann, I. Bahar, et al., The interface of protein structure, protein biophysics, and molecular evolution, Protein Science 21, 769 (2012).

[9] T. Tlusty, Self-referring DNA and protein: A remark on physical and geometrical aspects, Philosophical Transactions of the Royal Society A 374, 20150070 (2016).

[10] T. Tlusty, A. Libchaber, and J.-P. Eckmann, Physical model of the genotype-to-phenotype map of proteins, Physical Review X 7, 021037 (2017).

[11] S. Dutta, J.-P. Eckmann, A. Libchaber, and T. Tlusty, Green function of correlated genes in a minimal mechanical model of protein evolution, Proceedings of the National Academy of Sciences 115, E4559 (2018).

[12] J.-P. Eckmann, J. Rougemont, and T. Tlusty, Colloquium: Proteins: The physics of amorphous evolving matter, Reviews of Modern Physics 91, 031001 (2019).

[13] C. B. Anfinsen, Principles that govern the folding of protein chains, Science 181, 223 (1973).

[14] C. Chothia and A. M. Lesk, The relation between the divergence of sequence and structure in proteins., The EMBO journal 5, 823 (1986).

[15] J. Jumper, R. Evans, A. Pritzel, T. Green, M. Figurnov, O. Ronneberger, K. Tunyasuvunakool, R. Bates, A. Žídek Potapenko, et al., Highly accurate protein structure prediction with alphafold, nature 596, 583 (2021).

[16] M. Baek, F. DiMaio, I. Anishchenko, J. Dauparas, S. Ovchinnikov, G. R. Lee, J. Wang, Q. Cong, L. N. Kinch, R. D. Schaeffer, et al., Accurate prediction of protein structures and interactions using a three-track neural network, Science 373, 871 (2021).

[17] J. M. McBride, K. Polev, A. Abdirasulov, V. Reinharz, A. Grzybowski, and T. Tlusty, Alphafold2 can predict single-mutation effects, Physical Review Letters 131, 218401 (2023).

[18] J. M. McBride and T. Tlusty, Ai-predicted protein deformation encodes energy landscape perturbation, Physical Review Letters 133, 098401 (2024).

[19] J. Arroyo-Gomez, M. J. Murray, C. Guérillon, et al., Functional landscape of ubiquitin linkages couples k29-linked ubiquitylation to epigenome integrity, The EMBO Journal 44, 6944 (2025).

[20] E. J. van Aalst and B. J. Wylie, An in silico framework to visualize how cancer-associated mutations influence structural plasticity of the chemokine receptor CCR3, Protein Science 34, e70013 (2025).

[21] M. C. Lawrence and P. M. Colman, Shape complementarity at protein/protein interfaces (1993).

[22] A. Shulman-Peleg, R. Nussinov, and H. J. Wolfson, Recognition of functional sites in protein structures, Journal of molecular biology 339, 607 (2004).

[23] M. Chartier, E. Adriansen, and R. Najmanovich, Isomif finder: online detection of binding site molecular interaction field similarities, Bioinformatics 32, 621 (2016).

[24] S. Jones and J. M. Thornton, Analysis of protein-protein interaction sites using surface patches, Journal of molecular biology 272, 121 (1997).

[25] F. Glaser, T. Pupko, I. Paz, R. E. Bell, D. Bechor-Shental, E. Martz, and N. Ben-Tal, Consurf: identification of functional regions in proteins by surface-mapping of phylogenetic information, Bioinformatics 19, 163 (2003).

[26] F. M. Richards, Areas, volumes, packing, and protein structure, Annual review of biophysics and bioengineering 6, 151 (1977).

[27] F. B. Sheinerman, R. Norel, and B. Honig, Electrostatic aspects of protein–protein interactions, Current opinion in structural biology 10, 153 (2000).

[28] J. Kyte and R. F. Doolittle, A simple method for displaying the hydropathic character of a protein, Journal of molecular biology 157, 105 (1982).

[29] T. Kortemme, A. V. Morozov, and D. Baker, An orientation-dependent hydrogen bonding potential improves prediction of specificity and structure for proteins and protein–protein complexes, Journal of molecular biology 326, 1239 (2003).

[30] J. Konc and D. Janežič, Protein-protein binding-sites prediction by protein surface structure conservation, Journal of chemical information and modeling 47, 940 (2007).

[31] A. Kessel and N. Ben-Tal, Introduction to proteins: structure, function, and motion (Chapman and Hall/CRC, 2018).

[32] J. M. McBride, A. Koshevarnikov, M. Siek, B. A. Grzybowski, and T. Tlusty, Statistical survey of chemical and geometric patterns on protein surfaces as a blueprint for protein-mimicking nanoparticles, Small Structures 5, 2400086 (2024).

[33] B. Lee and F. M. Richards, The interpretation of protein structures: estimation of static accessibility, Journal of molecular biology 55, 379 (1971).

[34] M. L. Connolly, Analytical molecular surface calculation, Applied Crystallography 16, 548 (1983).

[35] D. La, J. Esquivel-Rodríguez, V. Venkatraman, B. Li, L. Sael, S. Ueng, S. Ahrendt, and D. Kihara, 3d-surfer: software for high-throughput protein surface comparison and analysis, Bioinformatics 25, 2843 (2009).

[36] S. Yin, E. A. Proctor, A. A. Lugovskoy, and N. V. Dokholyan, Fast screening of protein surfaces using geometric invariant fingerprints, Proceedings of the National Academy of Sciences 106, 16622 (2009).

[37] V. Venkatraman, Y. D. Yang, L. Sael, and D. Kihara, Protein-protein docking using region-based 3d zernike descriptors, BMC bioinformatics 10, 407 (2009).

[38] D. Kihara, L. Sael, R. Chikhi, and J. Esquivel-Rodriguez, Molecular surface representation using 3d zernike descriptors for protein shape comparison and docking, Current Protein and Peptide Science 12, 520 (2011).

[39] J. Hass and P. Koehl, How round is a protein? exploring protein structures for globularity using conformal mapping, Frontiers in molecular biosciences 1, 26 (2014).

[40] X. Zhu, Y. Xiong, and D. Kihara, Large-scale binding ligand prediction by improved patch-based method patch-surfer2. 0, Bioinformatics 31, 707 (2015).

[41] L. Sirugue, F. Langenfeld, N. Lagarde, and M. Montes, Plo3s: Protein local surficial similarity screening, Computational and Structural Biotechnology Journal 26, 1 (2024).

[42] H. Schweke, M.-H. Mucchielli, N. Chevrollier, S. Gosset, and A. Lopes, Surfmap: a software for mapping in two dimensions protein surface features, Journal of Chemical Information and Modeling 62, 1595 (2022).

[43] P. Gainza, F. Sverrisson, F. Monti, E. Rodola, D. Boscaini, M. M. Bronstein, and B. E. Correia, Deciphering interaction fingerprints from protein molecular surfaces using geometric deep learning, Nature methods 17, 184 (2020).

[44] M. Simonovsky and J. Meyers, Deeplytough: learning structural comparison of protein binding sites, Journal of chemical information and modeling 60, 2356 (2020).

[45] S. K. Mylonas, A. Axenopoulos, and P. Daras, Deepsurf: a surface-based deep learning approach for the prediction of ligand binding sites on proteins, Bioinformatics 37, 1681 (2021).

[46] S. Riahi, J. H. Lee, T. Sorenson, S. Wei, S. Jager, R. Olfati-Saber, Y. Zhou, A. Park, M. Wendt, H. Minoux, et al., Surface id: a geometry-aware system for protein molecular surface comparison, Bioinformatics 39, btad196 (2023).

[47] Y. Lin, L. Pan, Y. Li, Z. Liu, and X. Li, Exploiting hierarchical interactions for protein surface learning, IEEE Journal of Biomedical and Health Informatics 28, 1927 (2024).

[48] M. A. Fischler and R. C. Bolles, Random sample consensus: a paradigm for model fitting with applications to image analysis and automated cartography, Communications of the ACM 24, 381 (1981).

[49] P. J. Besl and N. D. McKay, Method for registration of 3-d shapes, in Sensor fusion IV: control paradigms and data structures, Vol. 1611 (Spie, 1992) pp. 586–606.

[50] Y. Chen and G. Medioni, Object modelling by registration of multiple range images, Image and vision computing 10, 145 (1992).

[51] B. Amberg, S. Romdhani, and T. Vetter, Optimal step nonrigid icp algorithms for surface registration, in 2007 IEEE conference on computer vision and pattern recognition (IEEE, 2007) pp. 1–8.

[52] A. Myronenko and X. Song, Point set registration: Coherent point drift, IEEE transactions on pattern analysis and machine intelligence 32, 2262 (2010).

[53] M. Ovsjanikov, M. Ben-Chen, J. Solomon, A. Butscher, and L. Guibas, Functional maps: a flexible representation of maps between shapes, ACM Transactions on Graphics (ToG) 31, 1 (2012).

[54] O. Van Kaick, H. Zhang, G. Hamarneh, and D. Cohen-Or, A survey on shape correspondence, in Computer graphics forum, Vol. 30 (Wiley Online Library, 2011) pp. 1681–1707.

[55] B. Deng, Y. Yao, R. M. Dyke, and J. Zhang, A survey of nonrigid 3d registration, in Computer Graphics Forum, Vol. 41 (Wiley Online Library, 2022) pp. 559–589.

[56] J. M. McBride, J.-P. Eckmann, and T. Tlusty, General theory of specific binding: insights from a genetic-mechanochemical protein model, Molecular Biology and Evolution 39, msac217 (2022).

[57] Y. Zhang and J. Skolnick, Tm-align: a protein structure alignment algorithm based on the tm-score, Nucleic acids research 33, 2302 (2005).

[58] F. P. Guengerich, Cytochrome p450 and chemical toxicology, Chemical research in toxicology 21, 70 (2008).

[59] T. Omura, Structural diversity of cytochrome p450 enzyme system, Journal of biochemistry 147, 297 (2010).

[60] Y. Vander Meersche, G. Cretin, A. Gheeraert, J.-C. Gelly, and T. Galochkina, Atlas: protein flexibility description from atomistic molecular dynamics simulations, Nucleic acids research 52, D384 (2024).

[61] H. M. Berman, J. Westbrook, Z. Feng, G. Gilliland, T. N. Bhat, H. Weissig, I. N. Shindyalov, and P. E. Bourne, The protein data bank, Nucleic acids research 28, 235 (2000).

[62] A. C. Y. Foo, P. M. Thompson, S.-H. Chen, R. Jadi, B. Lupo, E. F. DeRose, S. Arora, V. C. Placentra, L. Premkumar, Perera, L. C. Pedersen, N. Martin, and G. A. Mueller, The mosquito protein AEG12 displays both cytolytic and antiviral properties via a common lipid transfer mechanism, Proc. Natl. Acad. Sci. U.S.A. 118, 10.1073/pnas.2019251118 (2021).

[63] J.-L. Berry, Y. Xu, P. N. Ward, S. M. Lea, S. J. Matthews, and V. Pelicic, A Comparative Structure/Function Analysis of Two Type IV Pilin DNA Receptors Defines a Novel Mode of DNA Binding, Structure 24, 926 (2016).

[64] D. Lamb, A. W. Schüttelkopf, D. M. F. Van Aalten, and D. W. Brighty, Charge-Surrounded Pockets and Electrostatic Interactions with Small Ions Modulate the Activity of Retroviral Fusion Proteins, PLoS Pathog 7, e1001268 (2011).

[65] N. J. Byrne, A. C. Lee, J. Kostas, J. C. Reid, A. T. Partridge, S.-S. So, J. E. Cowan, P. Abeywickrema, H. Huang, M. Zebisch, J. J. Barker, S. M. Soisson, A. Brooun, and H.-P. Su, Development of a robust crystallization platform for immune receptor TREM2 using a crystallization chaperone strategy, Protein Expression and Purification 179, 105796 (2021).

[66] M. Steinegger and J. Söding, Mmseqs2 enables sensitive protein sequence searching for the analysis of massive data sets, Nature biotechnology 35, 1026 (2017).

[67] J. Lin, Divergence measures based on the shannon entropy, IEEE Transactions on Information theory 37, 145 (2002).

[68] M. Reuter, F.-E. Wolter, and N. Peinecke, Laplace–beltrami spectra as ‘shape-dna’of surfaces and solids, Computer-Aided Design 38, 342 (2006).

[69] P. Gainza, F. Sverrisson, F. Monti, E. Rodolà, D. Boscaini, M. Bronstein, and B. E. Correia, Deciphering interaction fingerprints from protein molecular surfaces using geometric deep learning, Nat Methods 17, 184 (2020).

[70] F. Mémoli, Gromov–wasserstein distances and the metric approach to object matching, Foundations of computational mathematics 11, 417 (2011).

[71] T. Vayer, L. Chapel, R. Flamary, R. Tavenard, and N. Courty, Fused gromov-wasserstein distance for structured objects: theoretical foundations and mathematical properties, arXiv preprint arXiv:1811.02834 (2018).

[72] R. Sinkhorn, Diagonal equivalence to matrices with prescribed row and column sums, The American Mathematical Monthly 74, 402 (1967).

[73] M. Cuturi, Sinkhorn distances: Lightspeed computation of optimal transport, Advances in neural information processing systems 26 (2013).

[74] M. R. Mitchell, T. Tlusty, and S. Leibler, Strain analysis of protein structures and low dimensionality of mechanical allosteric couplings, Proceedings of the National Academy of Sciences 113, E5847 (2016).

[75] R. Flamary, N. Courty, A. Gramfort, M. Z. Alaya, A. Boisbunon, S. Chambon, L. Chapel, A. Corenflos, K. Fatras, Fournier, L. Gautheron, N. T. Gayraud, H. Janati, A. Rakotomamonjy, I. Redko, A. Rolet, A. Schutz, V. Seguy, D. J. Sutherland, R. Tavenard, A. Tong, and T. Vayer, Pot: Python optimal transport, Journal of Machine Learning Research 22, 1 (2021).

[76] R. Flamary, C. Vincent-Cuaz, N. Courty, A. Gramfort, O. Kachaiev, H. Quang Tran, L. David, C. Bonet, N. Cassereau, T. Gnassounou, E. Tanguy, J. Delon, A. Collas, S. Mazelet, L. Chapel, T. Kerdoncuff, X. Yu, M. Feickert, P. Krzakala, T. Liu, and E. Fernandes Montesuma, Pot python optimal transport (version 0.9.5) (2024).

[77] J. Sun, M. Ovsjanikov, and L. Guibas, A concise and provably informative multi-scale signature based on heat diffusion, in Computer graphics forum, Vol. 28 (Wiley Online Library, 2009) pp. 1383–1392.

[78] M. Aubry, U. Schlickewei, and D. Cremers, The wave kernel signature: A quantum mechanical approach to shape analysis, in 2011 IEEE international conference on computer vision workshops (ICCV workshops) (IEEE, 2011) pp. 1626–1633.

[79] A. Raffo, U. Fugacci, S. Biasotti, and W. Rocchia, SHREC 2021: Protein surface dataset and ground truths (2021), accessed July 30, 2026.

