## Supplementary material for "Geometric–Chemical Distance Between Protein Surfaces": SI

### THE GENERAL IFACE FRAMEWORK

Let $S_\alpha$ and $S_\beta$ be two protein surfaces, each equipped with a collection of fields $F^\alpha = \{f_m^\alpha\}$ and $F^\beta = \{f_m^\beta\}$, where each $f_m^\alpha : S_\alpha \to \mathbb{R}^{n_m}$ and $f_m^\beta : S_\beta \to \mathbb{R}^{n_m}$ may represent a scalar field ($n_m = 1$), a vector field, or a tensor field. Although protein surfaces can be described more richly using vector and tensor fields (*e.g.*, electric fields, stress–strain[74], and polarizability tensors), they are still studied predominantly with scalar fields. Therefore, in this work, we restrict our analysis to **scalar** fields on protein surfaces—such as electrostatic potential, hydrogen bonding propensity, and hydrophobicity. Our goal is to define a distance metric $D(S_\alpha, S_\beta)$ that jointly captures both geometric and field-level differences.

**From Diffeomorphic Mapping to Probabilistic Soft Mapping for Field Transport.** Two surfaces are considered equivalent when there exists a diffeomorphism that maps one to the other while leaving all transported fields unchanged, *i.e.*,

$$S_\alpha \sim S_\beta \iff \exists\, \varphi \in \mathrm{Diff}(S_\alpha, S_\beta) \text{ such that } \varphi^* g_\beta = g_\alpha \quad \text{and} \quad \varphi^* F^\beta = F^\alpha.$$

Here, $\varphi^* F^\beta$ denotes the family of fields on $S_\alpha$ obtained by pulling back the fields in $F^\beta$, and $\varphi^* g_\beta$ denotes the metric on $S_\alpha$ obtained by pulling back $g_\beta$, via the diffeomorphism $\varphi\colon S_\alpha \to S_\beta$.

Equivalently, the similarity between the pullback fields and the native fields can be quantified using $L^p$ norms. We define the pullback distance between families of scalar fields as the $\ell^1$-sum of the componentwise $L^p$ distances:

$$d_F^{(p)}(F^\alpha, F^\beta; \varphi) := \sum_m \left( \int \left| f_\alpha^m - \varphi^* f_\beta^m \right|^p \, d\mu^\alpha \right)^{\frac{1}{p}}.$$

Here, $p \geq 1$ denotes the exponent of the $L^p$ norm. $d\mu^\alpha$ denotes the surface-area measure on the surface ($S_\alpha$) and it is given by $d\mu^\alpha \equiv \sqrt{|g_\alpha(x)|}\, d^2x$, where $|g_\alpha(x)|$ is the determinant of the metric tensor on the surface $S_\alpha$. Further, to compare the internal geometric structure, we compare the metric tensor fields using the pushforward map $\varphi_*$:

$$d_G^{(q)}(g_\alpha, g_\beta; \varphi) := \left( \int_{S_\beta} \left\| \varphi_* g_\alpha - g_\beta \right\|^q \, d\mu^\beta \right)^{\frac{1}{q}}.$$

Here $d\mu^\beta$ denotes the surface area measure on $S_\beta$, $q \geq 1$ is the exponent for the geometric discrepancy, and $\|\cdot\|$ denotes a pointwise tensor norm. The exponents $p$ and $q$ control the sensitivity of the field and geometric discrepancies, respectively. Furthermore, similar expressions can be obtained for the inverse mapping $\varphi^{-1}$. We can now define a distance measure, $D(S_\alpha, S_\beta)$, using the diffeomorphism mapping, as follows

$$\begin{aligned} D(S_\alpha, S_\beta) = \sum_m \zeta_m \Bigg( \frac{1}{2} \Big( \int_{S_\alpha} |f_\alpha^m - \varphi^* f_\beta^m|^p \, d\mu^\alpha \\ + \int_{S_\beta} |f_\beta^m - \varphi^{-1*} f_\alpha^m|^p \, d\mu^\beta \Big) \Bigg)^{\frac{1}{p}} \\ + \eta \Bigg( \frac{1}{2} \Big( \int_{S_\beta} \|\varphi_* g_\alpha - g_\beta\|^q \, d\mu^\beta \\ + \int_{S_\alpha} \|\varphi_*^{-1} g_\beta - g_\alpha\|^q \, d\mu^\alpha \Big) \Bigg)^{\frac{1}{q}}. \end{aligned} \tag{S1}$$

Here $\zeta_m$ and $\eta$ denote weighting coefficients for the $m^{\text{th}}$ scalar field and the metric tensor discrepancy, respectively. These weights control the relative importance of the terms and provide normalization so that the resulting quantities are dimensionless or measured in the same units. We note that the above metric (Eq. S1) is clearly symmetric under the exchange, $S_\alpha \leftrightarrow S_\beta$, and furthermore, it can be shown that it satisfies the properties of a metric.

**Probabilistic Soft Mapping.** We have described a distance metric that assumes the existence of a diffeomorphism between the surfaces. However, protein surfaces are often corrupted by noise and limited resolution and an *exact* diffeomorphism rarely exists, even between two conformers of the *same* protein. Therefore, rather than searching for a single hard map $\varphi : S_\alpha \to S_\beta$, we relax the correspondence to a probability distribution on the product space $S_\alpha \times S_\beta$. In other words, we allow each point $x \in S_\alpha$ to match several points $y \in S_\beta$ with varying likelihoods. In addition, this approach has the advantage that, instead of producing a continuous map, it can probabilistically map different parts of proteins that have similar patches. For example, two proteins may share only a pocket, while the other parts of the proteins differ from each other.

Formally, we introduce a coupling $p(x, y)$.

$$p(x, y) \in \mathcal{P}(S_\alpha \times S_\beta), \quad p(x, y) \geq 0,$$
$$\int_{S_\alpha} p(x, y) d\mu^\alpha(x) = \rho^\beta(y), \qquad \int_{S_\beta} p(x, y) d\mu^\beta(y) = \rho^\alpha(x),$$

where $\rho^\alpha(x)$ and $\rho^\beta(y)$ are the marginal distributions on the respective spaces. These marginal distributions represent, in a physical sense, the probability–or weight–with which each point on the surface should be considered when being mapped to the other. They can be chosen according to prior knowledge about which regions of one surface are more likely to be present and have a correspondence to regions on the other. They satisfy the normalization conditions:

$$\int_{S_\alpha} \rho^\alpha(x)\, d\mu^\alpha(x) = 1, \quad \int_{S_\beta} \rho^\beta(y)\, d\mu^\beta(y) = 1,$$

which ensure that $\rho^\alpha$ and $\rho^\beta$ are valid probability density functions on $S_\alpha$ and $S_\beta$, respectively. Incorporating these into the Eq. S1 leads to a distance which we call the soft distance:

$$D_{\text{soft}}(S_\alpha, S_\beta) = \sum_m \zeta_m \left( \frac{1}{2} \int_{S_\alpha}\int_{S_\beta} \left|f_\alpha^m(x) - f_\beta^m(y)\right|^p p(x,y)\, d\mu^\alpha(x)\, d\mu^\beta(y) + (S_\alpha \leftrightarrow S_\beta) \right)^{\frac{1}{p}} + \eta \left( \frac{1}{2} \int_{S_\alpha}\int_{S_\beta} \left\|g_\alpha(x) - g_\beta(y)\right\|^q p(x,y)\, d\mu^\alpha(x)\, d\mu^\beta(y) + (S_\alpha \leftrightarrow S_\beta) \right)^{\frac{1}{q}}.$$

We note that comparing metric tensors requires a transformation that involves the pushforward (*i.e.*, transporting) of tangent spaces between the surfaces. This, in turn, necessitates the introduction of additional structure into our framework and thereby increases complexity. To minimize this complexity and make the comparison more tractable, we relax the notion of local distance—originally encoded by the metric tensor—to a more global measure on the surface by using pairwise geodesic distances. Incorporating this into our formulation, the distance becomes

$$D^{\text{soft}}(S_\alpha, S_\beta) = D^{\text{soft}}_{\text{field}}(S_\alpha, S_\beta) + D^{\text{soft}}_{\text{geom}}(S_\alpha, S_\beta)\,, \quad \text{(S2)}$$

where

$$D^{\text{soft}}_{\text{field}}(S_\alpha, S_\beta) := \sum_m \zeta_m \Big( \frac{1}{2} \int_{S_\alpha}\int_{S_\beta} \left|f_\alpha^m(x) - f_\beta^m(y)\right|^p p(x,y)\, d\mu^\alpha(x)\, d\mu^\beta(y) + (S_\alpha \leftrightarrow S_\beta) \Big)^{\frac{1}{p}},$$

$$D^{\text{soft}}_{\text{geom}}(S_\alpha, S_\beta) = \eta \left( \int_{S_\alpha^2}\int_{S_\beta^2} \frac{1}{2} \left|d_\alpha(x,x') - d_\beta(y,y')\right|^q p(x,y)p(x',y') d\mu^\alpha(x)\, d\mu^\alpha(x')\, d\mu^\beta(y)\, d\mu^\beta(y') + (S_\alpha \leftrightarrow S_\beta) \right)^{\frac{1}{q}},$$

where $d_\alpha(x,x')$ and $d_\beta(y,y')$ denote the pairwise geodesic distances on $S_\alpha$ and $S_\beta$, respectively. Note that, since we are comparing pairs of points, we include an additional soft correspondence term $p(x', y')$ in the integral. Further, Eq. S2 is symmetric under exchange $S_\alpha \leftrightarrow S_\beta$. As we are going to use meshes, we add a term which further strengthens the global structure

$$h_S(x,x') = \int_\Omega k_S(x,z)\, k_S(z,x')\, \mathrm{d}z,$$

$$k_S(z,x') = \exp\left[ -\frac{1}{2} \left( \frac{d_S(z,x')}{\sigma_S} \right)^2 \right].$$

We finally define

$$G_S(x,x') = d_S(x,x') + \epsilon_g h_S(x,x'),$$

where $d_S(x,x')$ denotes the pairwise geodesic distance on the surface $S$, and $\epsilon_g$ and $\sigma_S$ are parameters controlling the contribution of the global structure. So our final structural/geometric term is defined as

$$D^{\text{soft}}_{\text{Structure}}(S_\alpha, S_\beta) = \eta \left( \int_{S_\alpha^2}\int_{S_\beta^2} \frac{1}{2} \left|G_\alpha(x,x') - G_\beta(y,y')\right|^q p(x,y)p(x',y')\, d\mu^\alpha(x)\, d\mu^\alpha(x')\, d\mu^\beta(y)\, d\mu^\beta(y') + (S_\alpha \leftrightarrow S_\beta) \right)^{\frac{1}{q}}.$$

**The IFACE Distance.** We now define our distance, referred to as the **IFACE distance**, using the soft-correspondence values, based on the top-$k$ normalized weights and their associated vertices. For each point, we select the $k(=2)$ largest values from the joint correspondence density weighted by local geometric measures:

$$\tilde{w}^{y_k}(x) \equiv \text{Normalize}\left( \text{Top}_k^y \left[ p(x,y) \cdot \sqrt{|g_\alpha(x)| \cdot |g_\beta(y)|} \right] \right),$$

$$\tilde{w}^{x_k}(y) \equiv \text{Normalize}\left( \text{Top}_k^x \left[ p(x,y) \cdot \sqrt{|g_\alpha(x)| \cdot |g_\beta(y)|} \right] \right).$$

Here, $p(x,y)$ denotes the soft correspondence *density* between points $x \in S_\alpha$ and $y \in S_\beta$, and $|g_\alpha(x)|$, $|g_\beta(y)|$ are the determinants of the metric tensors at $x$ and $y$, respectively. The `Top` operator selects the $k$ largest weighted densities for each fixed variable (either $x$ or $y$), and the `Normalize` operation ensures that the resulting weights form a valid probability distribution, *i.e.*, they integrate to 1 over the selected $k$-neighborhood.

Now, $\tilde{w}^{y_k}(x)$ represents a normalized soft distribution over the top-$k$ points $(y_k)$ on $S_\beta$ that most strongly correspond to a given point $x \in S_\alpha$, and vice versa for $\tilde{w}^{x_k}(y)$.

Then the IFACE distance is defined by the new weights in Eq. S3:

$$D_{\text{IFACE}}(S_\alpha, S_\beta) = \sum_m \zeta_m\, D^m_{\text{field}}(S_\alpha, S_\beta) + \eta\, D_{\text{structure}}(S_\alpha, S_\beta)\,, \quad \text{(S3)}$$

where

$$D^m_{\text{field}}(S_\alpha, S_\beta) = \frac{1}{2^{1/p}} \left( \sum_k \int_{S_\alpha} |f_\alpha^m(x) - f_\beta^m(y_k)|^p\, \tilde{w}^{y_k}(x)\, d^2x + \int_{S_\beta} |f_\alpha^m(x_k) - f_\beta^m(y)|^p\, \tilde{w}^{x_k}(y)\, d^2y \right)^{\frac{1}{p}}$$

$$D_{\text{structure}}(S_\alpha, S_\beta) = \frac{1}{2^{1/q}} \sum_{k,k'} \Bigg( \int_{S_\alpha^2} \|G_\alpha(x,x') - G_\beta(y_k, y'_{k'})\|^q \, \tilde{w}^{y_k}(x)\, \tilde{w}^{y_{k'}}(x')\, d^2x\, d^2x' + \int_{S_\beta^2} \|G_\alpha(x_k, x'_{k'}) - G_\beta(y,y')\|^q \, \tilde{w}^{x_k}(y)\, \tilde{w}^{x_{k'}}(y')\, d^2y\, d^2y' \Bigg)^{\frac{1}{q}}.$$

### DISCRETE PROTEIN SURFACE

We use the discrete form of the problem as described in the main text. Here, we describe the protein surface generation pipeline, calculation of the marginal distribution used in the method, and the optimization process.

**Protein Surface Generation.** We use a modified MaSIF pipeline (github.com/jomimc/masif_molecule) to generate protein surfaces comprising vertex positions, connectivity information, and feature files describing vertex-level attributes, including charge, hydrophobicity (Kyte–Doolittle), Gaussian curvature, mean curvature, amino acid identity, and the sequence index of the corresponding residue. The resulting protein meshes are preprocessed for analysis and contain a maximum of 3,000 vertices per mesh.

Because optimization becomes computationally expensive for large mesh sizes, we simplify protein surface meshes while consistently transferring per-vertex geometric and chemical properties to the reduced representation. Starting from an input surface mesh, we first perform a cleaning step to remove invalid elements and disconnected components, producing a watertight mesh. The cleaned mesh is then decimated using quadric edge collapse to obtain a target resolution, with the number of faces proportional to the desired vertex count. After decimation, small isolated components are removed, and the mesh is further refined using Laplacian smoothing to reduce discretization noise while preserving overall surface geometry.

To propagate vertex-wise properties from the original mesh to the simplified mesh, we establish a correspondence based on spatial proximity. Specifically, each vertex of the simplified mesh is mapped to its nearest vertex on the original mesh using a KD-tree search. For each mapped vertex, we define a local neighborhood on the original mesh using a fixed number of hops in the vertex adjacency graph. Continuous-valued properties are transferred by averaging over these neighborhoods, whereas discrete or categorical properties are directly inherited from the corresponding nearest vertices.

Finally, we compute geometric descriptors, including mean and Gaussian curvature, directly on the simplified mesh. The resulting meshes and associated properties are saved for downstream analysis, ensuring a consistent and computationally efficient surface representation with a bounded number of vertices.

**Algorithm S1: Protein Mesh Alignment and Coupling Matrix Estimation**

**Input:** Surface meshes $S_\alpha$ and $S_\beta$ with vertex sets $\{x_i\}, \{y_j\}$, associated vertex areas $\{\mu_i^\alpha\}, \{\mu_j^\beta\}$, and $m$-feature fields $f_i^{\alpha(m)}, f_j^{\beta(m)}$.

**Output:** Optimal coupling matrix $P_{ij}$.

1. Perform rigid alignment followed by feature-aware nonrigid alignment.
2. Transform vertices and features: $x_i \to \tilde{x}_i,\ f_i^{\alpha(m)} \to \tilde{f}_i^{\alpha(m)}$.
3. Construct descriptor vectors $\tilde{X}_i = [\tilde{x}_i, \tilde{f}_i^{\alpha(m)}], Y_j = [y_j, f_j^{\beta(m)}]$.
4. Compute Gaussian affinity kernel $A_{ij} = \exp(-\|\tilde{X}_i - Y_j\|^2/\delta^2)$ with $\delta = 0.01$.
5. Apply Sinkhorn normalization to the Gaussian affinity matrix.
6. Obtain initial coupling $P_{ij}$.
7. Optimize Eq. (S4) with $\lambda = 0.9$.
8. Update $P_{ij}$.
9. Re-optimize Eq. (S4) without entropic regularization.
10. Derive final coupling matrix $P_{ij}$.
11. Return $P_{ij}$.

**Calculation for marginal distributions.** In this section, we provide the procedure used to calculate marginal distribution for the protein surfaces. Let $\{\mu_i^\alpha\}$ and $\{\mu_j^\beta\}$ denote the vertex areas of surfaces $S_\alpha$ and $S_\beta$, respectively. Let $f_i^{\alpha m}$ and $f_j^{\beta m}$ denote the feature fields, where $m \in$ (Electrostatic Potential, Hydrophobicity, Hydrogen Bond Propensity, Mean curvature). We compute the marginal distributions $\rho_i^\alpha$ and $\rho_j^\beta$ as follows.

*Step 1: Area-based density (constant field).*

$$\rho_i^{\alpha(0)} = \frac{\mu_i^\alpha}{\sum_i \mu_i^\alpha}, \qquad \rho_j^{S_\beta(0)} = \frac{\mu_j^\beta}{\sum_j \mu_j^\beta}.$$

*Step 2: Global minimum feature value.*

$$f_{\min}^{(m)} = \min\left(\min_i f_i^{\alpha(m)},\ \min_j f_j^{\beta(m)}\right).$$

*Step 3: Area-weighted, shifted features.*

$$\tilde{f}_i^{\alpha(m)} = \left(f_i^{\alpha(m)} - f_{\min}^{(m)}\right)\mu_i^\alpha, \qquad \tilde{f}_j^{\beta(m)} = \left(f_j^{\beta(m)} - f_{\min}^{(m)}\right)\mu_j^\beta.$$

*Step 4: Feature normalization.*

$$\tilde{f}_i^{\alpha(m)} \leftarrow \frac{\tilde{f}_i^{\alpha(m)}}{\sum_i \tilde{f}_i^{\alpha(m)}}, \qquad \tilde{f}_j^{\beta(m)} \leftarrow \frac{\tilde{f}_j^{\beta(m)}}{\sum_j \tilde{f}_j^{\beta(m)}}.$$

*Step 5: Combine area and feature distributions.*

$$\rho_i^\alpha = \rho_i^{\alpha(0)} + \sum_m \tilde{f}_i^{\alpha(m)}, \qquad \rho_j^\beta = \rho_j^{\beta(0)} + \sum_m \tilde{f}_j^{\beta(m)}.$$

*Step 6: Final normalization.*

$$\rho_i^\alpha \leftarrow \frac{\rho_i^\alpha}{\sum_i \rho_i^\alpha}, \qquad \rho_j^\beta \leftarrow \frac{\rho_j^\beta}{\sum_j \rho_j^\beta}.$$

**Optimization for finding optimal coupling matrix.**

*Optimization.—* We perform a four-step refinement to obtain the coupling matrix $P_{ij}$, which includes mesh alignment followed by optimization steps. We first perform rigid alignment of the meshes using RANSAC[48] for coarse initialization, followed by Iterative Closest Point (ICP)[49,50] for refinement. After that, we use non-rigid alignment in the combined real and feature space, employing the coherent point drift algorithm[52]. We convert the point sets into a Gaussian kernel to obtain a similarity measure between the transformed and target points. Then, we convert the Gaussian kernel into a coupling/soft-correspondence matrix using the Sinkhorn update[72,73] with the marginal distribution. We feed that solution into the entropically regularized objective function for the field and structural terms with $\lambda = 0.9$ (this value is fixed across all examples). After that, we refine the coupling matrix by solving the objective function without entropic regularization. The algorithmic steps are given in Algorithm S1.

*Non-rigid alignment for initializing coupling matrix.—* After rigid alignment of meshes, we perform non-rigid registration in the joint geometric and feature space using the Coherent Point Drift (CPD) algorithm[52]. CPD models the source points as centroids of a Gaussian mixture and seeks a smooth non-rigid transformation that maximizes the likelihood of the target points under this model while enforcing motion coherence. Let $X \equiv [x_i, f_i^{\alpha(m)}]$ and $Y \equiv [y_j, f_j^{\beta(m)}]$ denote the source and target vertex sets, respectively, after rigid alignment in coordinate space, together with the $m$-feature-field values defined on those vertices. The non-rigid CPD transformation $T(\cdot)$ produces transformed points for source $\tilde{X} = \{T(X_i)\}$.

To quantify the similarity between transformed and target points, we construct a Gaussian kernel

$$A_{ij} = \exp\left(-\frac{\|\tilde{X}_i - Y_j\|^2}{\delta^2}\right),$$

which encodes pairwise affinities between $\tilde{X}$ and $Y$. We then interpret $A$ as an entropic-regularized similarity matrix and convert it into a soft correspondence (coupling) matrix by applying Sinkhorn iterations[72,73] to match prescribed marginal distributions. This yields a coupling matrix $P_{ij}$, which serves as the initial coupling (soft correspondence) matrix. We set $\delta = 0.01$ to enforce a tight similarity constraint. All steps are performed on unit-normalized features. For each feature field, joint min–max normalization is applied, where the scaling parameters are computed from the concatenation of the corresponding features and the values are mapped to $[-1, 1]$. Meshes are independently normalized to lie within the unit sphere.

*Entropic and entropic free regularized optimization of objective function.—* We use the initial coupling matrix $P_{ij}$ obtained using the above procedure as an initialization for the entropic optimization of the objective function combining the structural and field terms:

$$P_{ij}^{entro} = \arg\min_{P_{ij}} \Big[ (1-\lambda)\,\mathcal{F}(S_\alpha, S_\beta) + \lambda\,\mathcal{S}(S_\alpha, S_\beta) + \epsilon\, P_{ij} \log P_{ij} \Big],$$

$$\text{subject to} \sum_j P_{ij} = \rho_i^\alpha, \quad \sum_i P_{ij} = \rho_j^\beta, \qquad \text{(S4)}$$

where $\mathcal{F}(S_\alpha, S_\beta)$ and $\mathcal{S}(S_\alpha, S_\beta)$ are the field and structural terms defined in the Methods section of the main text for the optimization.We use unit-normalized meshes to compute the distance matrices used in the structural term.

*Weight parameters.—* The parameter $\lambda$ controls the balance between geometric correspondence and field alignment in the surface coupling matrix. We set $\lambda = 0.9$, giving dominant weight to the structural term so that the surface map remains geometrically continuous. The field term then acts as a secondary constraint that aligns regions with similar physicochemical fields. While this contribution can break geometric symmetries, the strong structural weighting preserves the global organization of the correspondence.

To verify that this choice achieves the intended balance, we examined a controlled toy system consisting of ellipsoidal surfaces. The surfaces were geometrically identical but carried different charge distributions, allowing the influence of the field term to be isolated from geometry. Across the explored range of $\lambda$, smaller values emphasize agreement of the field term, whereas larger values emphasize geometric consistency. We observe qualitatively stable behavior for $\lambda \gtrsim 0.85$ and therefore select $\lambda = 0.9$, which maintains structural continuity while allowing the field contribution to guide feature-consistent alignment across the surface.

*Entropic regularization parameter.—* The entropic regularization parameter $\epsilon$ is chosen adaptively to maintain numerical stability during optimization. Specifically,

$$\epsilon = \max\left(\frac{\max \mathcal{C}}{\kappa},\ \tau \cdot \mathrm{median}(\mathcal{C}),\ 10^{-6}\right),$$

where $\mathcal{C}$ denotes the collection of cost matrices used in the optimization, including the structural matrices $G_\alpha$, $G_\beta$ and the field-based cost term comparing aligned physicochemical features $\mathcal{F}(S_\alpha, S_\beta)$. The geometric and chemical terms are scaled to comparable magnitudes, so that $\lambda$ controls their relative contribution rather than compensating for differences in scale. The maximum and median are taken over all scalar entries of these matrices. The constant $\kappa$ prevents numerical underflow, while $\tau$ sets the regularization scale relative to the typical magnitude of the cost terms; we use $\kappa = 700$ and $\tau = 0.05$. This choice was found to be robust across datasets and does not affect the qualitative behavior of the distances.

The entropically regularized problem is first solved to obtain an initial coupling $P_{ij}^{\text{entro}}$. This solution is then used to initialize a second optimization without entropic regularization ($\epsilon = 0$), yielding the final optimal coupling matrix $P_{ij}$. All optimizations were performed using the POT Python library[75,76]. Across the dataset of 1080 protein pairs, the full pipeline—including optimization and distance computations—takes about 8.5 minutes per pair on average. Experiments were conducted on a workstation equipped with an Intel Core i9-14900K CPU (24 cores, 32 threads) and 64 GB RAM running Ubuntu 24.04.3 LTS. For rapid screening and large-scale comparison, meshes containing approximately 1,000 vertices substantially reduce the computational cost while preserving comparable classification and clustering performance in the present benchmarks. The feature-specific scaling relations described in Sec. (Normalization scale for distances) were used to transfer the fixed normalization ranges between mesh resolutions.

**Color transfer between the mapped surfaces.** Colors on the target surface $c_j^{\text{target}}$ were computed by aggregating probability-weighted colors from matched vertices on the source surface $c_i^{\text{source}}$ (e.g., target: `6XDS` and source: `6XRX` in Fig. 2), followed by normalization by the total incoming probability mass of the coupling matrix:

$$c_j^{\text{target}} = \frac{\sum_i P_{i\to j}\, c_i^{\text{source}}}{\sum_i P_{i\to j}}.$$

Here, the summation is restricted to the top two matches per $i$.

### COMMENTS ON GEOMETRY-BASED CORRESPONDENCE METHODS

We assessed geometry-driven correspondence methods on protein surfaces using a representative functional correspondence approach[53], widely used in shape matching, as a benchmark. These methods did not consistently distinguish structural divergence from conformational variability on the dataset considered in the main text. For brevity, we do not present quantitative benchmarks here.

These observations suggest that the assumptions underlying such methods—particularly the reliance on near-isometric structure for stable spectral representations—may be frequently violated for protein surfaces in a functionally meaningful way. Furthermore, the geometric descriptors (e.g., Heat Kernel Signature[77] and Wave Kernel Signature[78]) employed in these approaches may be insufficient to capture the high geometric complexity and roughness characteristic of protein surfaces, thereby limiting their effectiveness in this setting.

Taken together, these observations highlight the limitations of purely geometric approaches and motivate the joint geometric–chemical formulation introduced in IFACE.

### NORMALIZATION SCALE FOR DISTANCES

To avoid recomputing the normalization ranges from each evaluation dataset, we constructed a fixed reference set containing 130 protein surfaces. Of these, 65 were selected from the test partition of the SHREC 2021 protein-surface benchmark by choosing one representative surface from each of its 65 PDB-based classes[79]. In the SHREC PDB-based classification, each class contains the available NMR conformations associated with the same PDB entry. Selecting one surface per class therefore avoids overrepresenting proteins for which a larger number of conformations is available. The remaining 65 surfaces were drawn from the family and conformer collections considered in this study.

The complete reference set generates 8,385 distinct unordered, non-self protein-pair comparisons. Because these pairs span proteins drawn from several structural classes and datasets, they produce broad distributions of structural and physicochemical distances. The resulting extrema therefore provide a broad empirical normalization scale for the present analysis. These extrema should, however, be interpreted as empirical reference bounds rather than absolute theoretical bounds on the corresponding distances.

TABLE S1. Descriptor-specific approximate scaling exponents and fixed normalization ranges transferred from 1,000 to 3,000 vertices.

| Distance | $p_f$ | Reference minimum | Reference maximum |
|---|---|---|---|
| Structural | 0.9 | 3398 | 104856 |
| Mean curvature | 1.3 | 490 | 2038 |
| Hydrogen bond | 1.3 | 225 | 2008 |
| Hydrophobicity | 1.3 | 3367 | 35990 |
| Electrostatics | 1.2 | 1220 | 125866 |

All reference surfaces were sampled at $N_0 = 1000$ vertices to reduce the computational cost of evaluating the 8,385 pairwise comparisons. For each distance component $f$, the global minimum and maximum were computed over the reference set.

The reference ranges were transferred to the 3000-vertex benchmark resolution using the descriptor-specific scaling relation

$$d_f(N) = d_f(N_0)\left(\frac{N}{N_0}\right)^{p_f},$$

where $p_f$ is the scaling exponent for component $f$. For each of the $K = 1080$ protein pairs evaluated at both resolutions,

$$p_{f,k} = \frac{\log\left[d_{f,k}(3000)/d_{f,k}(1000)\right]}{\log(3000/1000)},$$

and the final exponent was obtained as

$$p_f = \frac{1}{K}\sum_{k=1}^{K} p_{f,k}.$$

The fitted exponents were used to scale the reference extrema from 1000 to 3000 vertices,

$$d_{f,\text{min/max}}(3000) = d_{f,\text{min/max}}(1000)\, 3^{p_f}.$$

The descriptor-specific scaling exponents and fixed normalization ranges are summarized in Table S1.

Each raw distance was then scaled by the corresponding reference range,

$$\bar{d}_f = \frac{d_f}{d_{f,\max} - d_{f,\min}}.$$

The reference minimum is not subtracted such that zero raw distance remains zero. The same fixed ranges were used for all family and conformer benchmarks, avoiding benchmark-specific rescaling while allowing the normalization scale to be transferred across mesh resolutions.

### FULL ABLATION STUDY

Complete conformer- and family-benchmark feature-combination results are reported in Tables S2 and S3. Complete family-level clustering results are reported in Tables S4 and S5; leave-one-feature-out IFACE results are reported in Tables S6 and S7; and the corresponding JSD ablations are reported in Tables S8 and S9.

The directed and bidirectional threshold success percentages are reported in Tables S10 and S11.

TABLE S2. Ablation study of correspondence-based and distribution-based (JSD) feature combinations for binary classification of inter- and intra-protein protein pairs for conformer benchmark. For each correspondence-based feature combination, the structural distance is paired with its distribution-based counterpart, the JSD Gaussian curvature distance.

| Feature Combination | Correspondence-based | | Distribution-based (JSD) | |
|---|---|---|---|---|
| | AP | ROC-AUC | AP | ROC-AUC |
| Electrostatic | 0.9736 | 0.9879 | 0.9501 | 0.9847 |
| HBond | 0.9226 | 0.9581 | 0.3894 | 0.7018 |
| Hydrophobicity | 0.9013 | 0.9513 | 0.7314 | 0.8996 |
| Mean Curvature | 0.8107 | 0.8926 | 0.3299 | 0.6192 |
| Structural | 0.8939 | 0.9166 | 0.2658 | 0.5745 |
| Electrostatic + Hydrophobicity | 0.9452 | 0.9734 | 0.9933 | 0.9977 |
| HBond + Electrostatic | 0.9673 | 0.9817 | 0.9533 | 0.9861 |
| HBond + Hydrophobicity | 0.9215 | 0.9584 | 0.7629 | 0.9072 |
| Mean Curvature + Electrostatic | 0.8618 | 0.9298 | 0.9302 | 0.9784 |
| Mean Curvature + HBond | 0.8568 | 0.9246 | 0.4029 | 0.7009 |
| Mean Curvature + Hydrophobicity | 0.8822 | 0.9390 | 0.7108 | 0.8820 |
| Structural + Electrostatic | 0.9111 | 0.9274 | 0.9386 | 0.9821 |
| Structural + HBond | 0.9138 | 0.9274 | 0.4007 | 0.7007 |
| Structural + Hydrophobicity | 0.9205 | 0.9349 | 0.7307 | 0.8875 |
| Structural + Mean Curvature | 0.9118 | 0.9281 | 0.3210 | 0.6299 |
| HBond + Electrostatic + Hydrophobicity | 0.9490 | 0.9737 | 0.9929 | 0.9975 |
| Mean Curvature + Electrostatic + Hydrophobicity | 0.9111 | 0.9550 | 0.9868 | 0.9951 |
| Mean Curvature + HBond + Electrostatic | 0.8928 | 0.9468 | 0.9270 | 0.9787 |
| Mean Curvature + HBond + Hydrophobicity | 0.8970 | 0.9467 | 0.7252 | 0.8862 |
| Structural + Electrostatic + Hydrophobicity | 0.9265 | 0.9418 | 0.9923 | 0.9974 |
| Structural + HBond + Electrostatic | 0.9221 | 0.9355 | 0.9470 | 0.9841 |
| Structural + HBond + Hydrophobicity | 0.9240 | 0.9409 | 0.7579 | 0.8964 |
| Structural + Mean Curvature + Electrostatic | 0.9209 | 0.9363 | 0.9185 | 0.9756 |
| Structural + Mean Curvature + HBond | 0.9186 | 0.9350 | 0.4102 | 0.7026 |
| Structural + Mean Curvature + Hydrophobicity | 0.9205 | 0.9402 | 0.6997 | 0.8692 |
| Mean Curvature + HBond + Electrostatic + Hydrophobicity | 0.9202 | 0.9592 | 0.9863 | 0.9949 |
| Structural + HBond + Electrostatic + Hydrophobicity | 0.9287 | 0.9467 | 0.9919 | 0.9972 |
| Structural + Mean Curvature + Electrostatic + Hydrophobicity | 0.9269 | 0.9463 | 0.9848 | 0.9946 |
| Structural + Mean Curvature + HBond + Electrostatic | 0.9255 | 0.9424 | 0.9202 | 0.9763 |
| Structural + Mean Curvature + HBond + Hydrophobicity | 0.9231 | 0.9444 | 0.7182 | 0.8762 |
| All Features | 0.9284 | 0.9496 | 0.9837 | 0.9942 |

TABLE S3. Ablation study of correspondence-based and distribution-based (JSD) feature combinations for binary classification of inter- and intra-family protein pairs. For each correspondence-based feature combination, the structural distance is paired with its distribution-based counterpart, the JSD Gaussian curvature distance.

| Feature Combination | Correspondence-based | | Distribution-based (JSD) | |
|---|---|---|---|---|
| | AP | ROC-AUC | AP | ROC-AUC |
| Electrostatic | 0.4650 | 0.7411 | 0.3582 | 0.7128 |
| HBond | 0.6402 | 0.8068 | 0.3556 | 0.7951 |
| Hydrophobicity | 0.5874 | 0.8617 | 0.5019 | 0.8277 |
| Mean Curvature | 0.8013 | 0.8985 | 0.7003 | 0.9336 |
| Structural | 0.4787 | 0.8371 | 0.6134 | 0.9056 |
| Electrostatic + Hydrophobicity | 0.5919 | 0.8659 | 0.4720 | 0.8131 |
| HBond + Electrostatic | 0.6535 | 0.8107 | 0.4591 | 0.7950 |
| HBond + Hydrophobicity | 0.6252 | 0.8359 | 0.5007 | 0.8323 |
| Mean Curvature + Electrostatic | 0.8058 | 0.9061 | 0.4688 | 0.8004 |
| Mean Curvature + HBond | 0.8108 | 0.8748 | 0.7321 | 0.9038 |
| Mean Curvature + Hydrophobicity | 0.8277 | 0.9215 | 0.7665 | 0.8990 |
| Structural + Electrostatic | 0.5023 | 0.8395 | 0.4659 | 0.8004 |
| Structural + HBond | 0.7290 | 0.9342 | 0.7787 | 0.9162 |
| Structural + Hydrophobicity | 0.7197 | 0.9260 | 0.7977 | 0.9024 |
| Structural + Mean Curvature | 0.7313 | 0.9514 | 0.6981 | 0.9320 |
| HBond + Electrostatic + Hydrophobicity | 0.6279 | 0.8399 | 0.4961 | 0.8266 |
| Mean Curvature + Electrostatic + Hydrophobicity | 0.8301 | 0.9261 | 0.5696 | 0.8483 |
| Mean Curvature + HBond + Electrostatic | 0.8168 | 0.8803 | 0.5613 | 0.8357 |
| Mean Curvature + HBond + Hydrophobicity | 0.8042 | 0.8983 | 0.7209 | 0.8905 |
| Structural + Electrostatic + Hydrophobicity | 0.7157 | 0.9247 | 0.5938 | 0.8553 |
| Structural + HBond + Electrostatic | 0.7210 | 0.9318 | 0.5796 | 0.8432 |
| Structural + HBond + Hydrophobicity | 0.8469 | 0.9562 | 0.7697 | 0.9014 |
| Structural + Mean Curvature + Electrostatic | 0.7316 | 0.9520 | 0.5190 | 0.8481 |
| Structural + Mean Curvature + HBond | 0.9434 | 0.9838 | 0.8241 | 0.9299 |
| Structural + Mean Curvature + Hydrophobicity | 0.8556 | 0.9736 | 0.8526 | 0.9241 |
| Mean Curvature + HBond + Electrostatic + Hydrophobicity | 0.8070 | 0.9014 | 0.5787 | 0.8560 |
| Structural + HBond + Electrostatic + Hydrophobicity | 0.8397 | 0.9543 | 0.6138 | 0.8646 |
| Structural + Mean Curvature + Electrostatic + Hydrophobicity | 0.8523 | 0.9730 | 0.6606 | 0.8792 |
| Structural + Mean Curvature + HBond + Electrostatic | 0.9401 | 0.9826 | 0.6335 | 0.8687 |
| Structural + Mean Curvature + HBond + Hydrophobicity | 0.9513 | 0.9835 | 0.8429 | 0.9231 |
| All Features | 0.9487 | 0.9829 | 0.6799 | 0.8866 |

TABLE S4. Adjusted Rand Index (ARI) for family-level clustering using all clustering algorithms and distance measures. Higher values indicate better agreement between predicted clusters and the ground-truth protein family labels.

| **Algorithm** | **IFACE (All Features)** | **JSD All Features** | **LB Distance** | **MaSIF Mean** | **MaSIF Mean+Max** | **SurfaceID Mean** | **SurfaceID Mean+Max** |
|---|---|---|---|---|---|---|---|
| Single | 0.653 | 0.300 | 0.188 | 0.537 | 0.417 | 0.205 | 0.153 |
| Complete | 0.899 | 0.556 | 0.424 | 0.578 | 0.380 | 0.218 | 0.328 |
| Average | 0.889 | 0.358 | 0.473 | 0.631 | 0.408 | 0.249 | 0.114 |
| Weighted | 0.889 | 0.358 | 0.513 | 0.578 | 0.445 | 0.262 | 0.328 |
| CMDS(4D)+K-means | 0.739 | 0.476 | 0.512 | 0.681 | 0.566 | 0.358 | 0.328 |
| CMDS(4D)+Ward | 0.739 | 0.451 | 0.512 | 0.713 | 0.558 | 0.249 | 0.439 |
| CMDS(5D)+K-means | 0.899 | 0.571 | 0.580 | 0.631 | 0.697 | 0.358 | 0.399 |
| CMDS(5D)+Ward | 0.899 | 0.571 | 0.437 | 0.631 | 0.697 | 0.249 | 0.281 |
| CMDS(6D)+K-means | 0.899 | 0.565 | 0.512 | 0.578 | 0.697 | 0.358 | 0.255 |
| CMDS(6D)+Ward | 0.899 | 0.495 | 0.437 | 0.578 | 0.697 | 0.249 | 0.242 |
| CMDS(8D)+K-means | 1.000 | 0.392 | 0.682 | 0.631 | 0.776 | 0.358 | 0.308 |
| CMDS(8D)+Ward | 1.000 | 0.357 | 0.748 | 0.631 | 0.697 | 0.249 | 0.328 |
| CMDS(10D)+K-means | 0.936 | 0.502 | 0.812 | 0.631 | 0.697 | 0.358 | 0.190 |
| CMDS(10D)+Ward | 0.949 | 0.502 | 0.748 | 0.631 | 0.566 | 0.249 | 0.328 |
| CMDS(15D)+K-means | 0.899 | 0.357 | 0.682 | 0.631 | 0.566 | 0.358 | 0.255 |
| CMDS(15D)+Ward | 0.899 | 0.502 | 0.748 | 0.631 | 0.493 | 0.249 | 0.328 |
| CMDS(20D)+K-means | 0.923 | 0.412 | 0.512 | 0.631 | 0.512 | 0.358 | 0.255 |
| CMDS(20D)+Ward | 0.899 | 0.357 | 0.748 | 0.631 | 0.697 | 0.249 | 0.328 |
| CMDS(80%)+K-means | 0.655 | 0.309 | 0.369 | 0.586 | 0.566 | 0.164 | 0.328 |
| CMDS(80%)+Ward | 0.936 | 0.309 | 0.412 | 0.586 | 0.697 | 0.127 | 0.439 |
| CMDS(90%)+K-means | 0.899 | 0.392 | 0.512 | 0.627 | 0.512 | 0.164 | 0.255 |
| CMDS(90%)+Ward | 0.923 | 0.357 | 0.512 | 0.618 | 0.697 | 0.127 | 0.242 |
| CMDS(95%)+K-means | 0.655 | 0.357 | 0.512 | 0.631 | 0.512 | 0.358 | 0.308 |
| CMDS(95%)+Ward | 0.899 | 0.502 | 0.437 | 0.631 | 0.493 | 0.249 | 0.328 |
| CMDS(99%)+K-means | 0.691 | 0.369 | 0.682 | 0.631 | 0.439 | 0.358 | 0.255 |
| CMDS(99%)+Ward | 0.899 | 0.357 | 0.748 | 0.631 | 0.697 | 0.249 | 0.328 |

TABLE S5. Cluster purity obtained using all clustering algorithms for each distance measure on the family benchmark. Higher values indicate better agreement between predicted clusters and the ground-truth protein family labels.

| **Algorithm** | **IFACE (All Features)** | **JSD All Features** | **LB Distance** | **MaSIF Mean** | **MaSIF Mean+Max** | **SurfaceID Mean** | **SurfaceID Mean+Max** |
|---|---|---|---|---|---|---|---|
| Single | 0.800 | 0.640 | 0.600 | 0.840 | 0.680 | 0.600 | 0.560 |
| Complete | 0.920 | 0.720 | 0.760 | 0.840 | 0.760 | 0.640 | 0.640 |
| Average | 0.920 | 0.680 | 0.760 | 0.840 | 0.720 | 0.680 | 0.560 |
| Weighted | 0.920 | 0.680 | 0.760 | 0.840 | 0.720 | 0.680 | 0.640 |
| CMDS(4D)+K-means | 0.920 | 0.720 | 0.800 | 0.880 | 0.800 | 0.720 | 0.640 |
| CMDS(4D)+Ward | 0.920 | 0.720 | 0.800 | 0.920 | 0.800 | 0.680 | 0.720 |
| CMDS(5D)+K-means | 0.920 | 0.760 | 0.840 | 0.840 | 0.840 | 0.720 | 0.640 |
| CMDS(5D)+Ward | 0.920 | 0.760 | 0.760 | 0.840 | 0.840 | 0.680 | 0.680 |
| CMDS(6D)+K-means | 0.920 | 0.720 | 0.800 | 0.840 | 0.840 | 0.720 | 0.640 |
| CMDS(6D)+Ward | 0.920 | 0.680 | 0.760 | 0.840 | 0.840 | 0.680 | 0.640 |
| CMDS(8D)+K-means | 1.000 | 0.720 | 0.880 | 0.840 | 0.880 | 0.720 | 0.640 |
| CMDS(8D)+Ward | 1.000 | 0.680 | 0.880 | 0.840 | 0.840 | 0.680 | 0.640 |
| CMDS(10D)+K-means | 0.960 | 0.800 | 0.920 | 0.840 | 0.840 | 0.720 | 0.600 |
| CMDS(10D)+Ward | 0.960 | 0.800 | 0.880 | 0.840 | 0.800 | 0.680 | 0.640 |
| CMDS(15D)+K-means | 0.920 | 0.680 | 0.880 | 0.840 | 0.800 | 0.720 | 0.640 |
| CMDS(15D)+Ward | 0.920 | 0.800 | 0.880 | 0.840 | 0.800 | 0.680 | 0.640 |
| CMDS(20D)+K-means | 0.960 | 0.720 | 0.800 | 0.840 | 0.760 | 0.720 | 0.640 |
| CMDS(20D)+Ward | 0.920 | 0.680 | 0.880 | 0.840 | 0.840 | 0.680 | 0.640 |
| CMDS(80%)+K-means | 0.920 | 0.680 | 0.720 | 0.840 | 0.800 | 0.600 | 0.640 |
| CMDS(80%)+Ward | 0.960 | 0.680 | 0.760 | 0.840 | 0.840 | 0.560 | 0.720 |
| CMDS(90%)+K-means | 0.920 | 0.720 | 0.800 | 0.840 | 0.760 | 0.600 | 0.640 |
| CMDS(90%)+Ward | 0.960 | 0.680 | 0.800 | 0.840 | 0.840 | 0.560 | 0.640 |
| CMDS(95%)+K-means | 0.920 | 0.680 | 0.800 | 0.840 | 0.760 | 0.720 | 0.640 |
| CMDS(95%)+Ward | 0.920 | 0.800 | 0.760 | 0.840 | 0.800 | 0.680 | 0.640 |
| CMDS(99%)+K-means | 0.920 | 0.680 | 0.880 | 0.840 | 0.760 | 0.720 | 0.640 |
| CMDS(99%)+Ward | 0.920 | 0.680 | 0.880 | 0.840 | 0.840 | 0.680 | 0.640 |

TABLE S6. Leave-one-feature-out clustering ablation for IFACE on the family benchmark. Entries report Adjusted Rand Index (ARI) across hierarchical clustering and CMDS-based clustering variants.

| Algorithm | All Features | w/o Structural | w/o Mean Curvature | w/o Hydrogen Bond | w/o Electrostatics | w/o Hydrophobicity | Chemical Only | Geometry Only |
|---|---|---|---|---|---|---|---|---|
| Single | 0.653 | 0.327 | 0.327 | 0.867 | 0.889 | 0.889 | 0.256 | 0.669 |
| Complete | 0.899 | 0.726 | 0.824 | 0.867 | 0.899 | 0.936 | 0.264 | 0.867 |
| Average | 0.889 | 0.388 | 0.699 | 0.867 | 0.889 | 0.899 | 0.388 | 0.867 |
| Weighted | 0.889 | 0.559 | 0.699 | 0.867 | 0.889 | 0.936 | 0.323 | 0.867 |
| CMDS(4D)+K-means | 0.739 | 0.766 | 0.675 | 0.739 | 0.739 | 0.675 | 0.264 | 0.675 |
| CMDS(4D)+Ward | 0.739 | 0.756 | 0.675 | 0.739 | 0.739 | 0.675 | 0.213 | 0.675 |
| CMDS(5D)+K-means | 0.899 | 0.698 | 0.777 | 0.691 | 0.645 | 0.675 | 0.328 | 0.675 |
| CMDS(5D)+Ward | 0.899 | 0.698 | 0.840 | 0.691 | 0.899 | 0.691 | 0.540 | 0.691 |
| CMDS(6D)+K-means | 0.899 | 0.791 | 0.521 | 0.691 | 0.899 | 0.899 | 0.328 | 0.691 |
| CMDS(6D)+Ward | 0.899 | 0.791 | 0.485 | 0.691 | 0.899 | 0.899 | 0.540 | 0.691 |
| CMDS(8D)+K-means | 1.000 | 0.949 | 0.651 | 0.691 | 1.000 | 0.936 | 0.264 | 0.691 |
| CMDS(8D)+Ward | 1.000 | 0.911 | 0.912 | 0.660 | 1.000 | 0.899 | 0.540 | 0.691 |
| CMDS(10D)+K-means | 0.936 | 0.924 | 0.854 | 0.739 | 0.936 | 0.936 | 0.232 | 0.582 |
| CMDS(10D)+Ward | 0.949 | 0.849 | 0.854 | 0.739 | 0.949 | 0.936 | 0.540 | 0.691 |
| CMDS(15D)+K-means | 0.899 | 0.572 | 0.593 | 0.673 | 0.899 | 0.899 | 0.264 | 0.582 |
| CMDS(15D)+Ward | 0.899 | 0.923 | 0.811 | 0.691 | 0.923 | 0.936 | 0.559 | 0.691 |
| CMDS(20D)+K-means | 0.923 | 0.513 | 0.730 | 0.675 | 0.923 | 0.899 | 0.414 | 0.691 |
| CMDS(20D)+Ward | 0.899 | 0.923 | 0.854 | 0.675 | 1.000 | 0.936 | 0.328 | 0.675 |
| CMDS(80%)+K-means | 0.655 | 0.584 | 0.854 | 0.675 | 0.936 | 0.936 | 0.362 | 0.739 |
| CMDS(80%)+Ward | 0.936 | 0.605 | 0.854 | 0.675 | 1.000 | 0.936 | 0.421 | 0.867 |
| CMDS(90%)+K-means | 0.899 | 0.572 | 0.924 | 0.673 | 0.899 | 0.899 | 0.264 | 0.582 |
| CMDS(90%)+Ward | 0.923 | 0.846 | 0.811 | 0.691 | 0.923 | 0.936 | 0.559 | 0.691 |
| CMDS(95%)+K-means | 0.655 | 0.539 | 0.851 | 0.691 | 0.897 | 0.899 | 0.362 | 0.691 |
| CMDS(95%)+Ward | 0.899 | 0.923 | 0.811 | 0.675 | 1.000 | 0.936 | 0.328 | 0.691 |
| CMDS(99%)+K-means | 0.691 | 0.599 | 0.730 | 0.691 | 0.923 | 0.899 | 0.264 | 0.691 |
| CMDS(99%)+Ward | 0.899 | 0.605 | 0.854 | 0.675 | 1.000 | 0.936 | 0.328 | 0.675 |

TABLE S7. Leave-one-feature-out clustering ablation for IFACE on the family benchmark. Entries report cluster purity across hierarchical clustering and CMDS-based clustering variants.

| Algorithm | All Features | w/o Structural | w/o Mean Curvature | w/o Hydrogen Bond | w/o Electrostatics | w/o Hydrophobicity | Chemical Only | Geometry Only |
|---|---|---|---|---|---|---|---|---|
| Single | 0.800 | 0.680 | 0.680 | 0.920 | 0.920 | 0.920 | 0.640 | 0.800 |
| Complete | 0.920 | 0.880 | 0.880 | 0.920 | 0.920 | 0.960 | 0.680 | 0.920 |
| Average | 0.920 | 0.680 | 0.800 | 0.920 | 0.920 | 0.920 | 0.680 | 0.920 |
| Weighted | 0.920 | 0.760 | 0.800 | 0.920 | 0.920 | 0.960 | 0.720 | 0.920 |
| CMDS(4D)+K-means | 0.920 | 0.880 | 0.920 | 0.920 | 0.920 | 0.920 | 0.640 | 0.920 |
| CMDS(4D)+Ward | 0.920 | 0.880 | 0.920 | 0.920 | 0.920 | 0.920 | 0.640 | 0.920 |
| CMDS(5D)+K-means | 0.920 | 0.880 | 0.840 | 0.920 | 0.880 | 0.920 | 0.680 | 0.920 |
| CMDS(5D)+Ward | 0.920 | 0.880 | 0.880 | 0.920 | 0.920 | 0.920 | 0.760 | 0.920 |
| CMDS(6D)+K-means | 0.920 | 0.920 | 0.800 | 0.920 | 0.920 | 0.920 | 0.680 | 0.920 |
| CMDS(6D)+Ward | 0.920 | 0.920 | 0.760 | 0.920 | 0.920 | 0.920 | 0.760 | 0.920 |
| CMDS(8D)+K-means | 1.000 | 0.960 | 0.920 | 0.920 | 1.000 | 0.960 | 0.640 | 0.920 |
| CMDS(8D)+Ward | 1.000 | 0.960 | 0.960 | 0.880 | 1.000 | 0.920 | 0.760 | 0.920 |
| CMDS(10D)+K-means | 0.960 | 0.960 | 0.920 | 0.920 | 0.960 | 0.960 | 0.640 | 0.880 |
| CMDS(10D)+Ward | 0.960 | 0.920 | 0.920 | 0.920 | 0.960 | 0.960 | 0.760 | 0.920 |
| CMDS(15D)+K-means | 0.920 | 0.760 | 0.880 | 0.920 | 0.920 | 0.920 | 0.640 | 0.880 |
| CMDS(15D)+Ward | 0.920 | 0.960 | 0.880 | 0.920 | 0.960 | 0.960 | 0.760 | 0.920 |
| CMDS(20D)+K-means | 0.960 | 0.760 | 0.840 | 0.920 | 0.960 | 0.920 | 0.680 | 0.920 |
| CMDS(20D)+Ward | 0.920 | 0.960 | 0.920 | 0.920 | 1.000 | 0.960 | 0.680 | 0.920 |
| CMDS(80%)+K-means | 0.920 | 0.840 | 0.920 | 0.920 | 0.960 | 0.960 | 0.720 | 0.920 |
| CMDS(80%)+Ward | 0.960 | 0.800 | 0.920 | 0.920 | 1.000 | 0.960 | 0.720 | 0.920 |
| CMDS(90%)+K-means | 0.920 | 0.760 | 0.960 | 0.920 | 0.920 | 0.920 | 0.640 | 0.880 |
| CMDS(90%)+Ward | 0.960 | 0.920 | 0.880 | 0.920 | 0.960 | 0.960 | 0.760 | 0.920 |
| CMDS(95%)+K-means | 0.920 | 0.800 | 0.920 | 0.920 | 0.920 | 0.920 | 0.720 | 0.920 |
| CMDS(95%)+Ward | 0.920 | 0.960 | 0.880 | 0.920 | 1.000 | 0.960 | 0.680 | 0.920 |
| CMDS(99%)+K-means | 0.920 | 0.800 | 0.840 | 0.920 | 0.960 | 0.920 | 0.640 | 0.920 |
| CMDS(99%)+Ward | 0.920 | 0.800 | 0.920 | 0.920 | 1.000 | 0.960 | 0.680 | 0.920 |

TABLE S8. Leave-one-feature-out clustering ablation for matched JSD distances on the family benchmark. Entries report Adjusted Rand Index (ARI) across hierarchical clustering and CMDS-based clustering variants.

| Algorithm | All Features | w/o Structural | w/o Mean Curvature | w/o Hydrogen Bond | w/o Electrostatics | w/o Hydrophobicity | Chemical Only | Geometry Only |
|---|---|---|---|---|---|---|---|---|
| Single | 0.300 | 0.282 | 0.282 | 0.282 | 0.335 | 0.461 | 0.282 | 0.649 |
| Complete | 0.556 | 0.495 | 0.248 | 0.493 | 0.722 | 0.357 | 0.286 | 0.756 |
| Average | 0.358 | 0.280 | 0.280 | 0.637 | 0.727 | 0.469 | 0.286 | 0.766 |
| Weighted | 0.358 | 0.280 | 0.286 | 0.580 | 0.760 | 0.469 | 0.286 | 0.766 |
| CMDS(4D)+K-means | 0.476 | 0.235 | 0.216 | 0.456 | 0.494 | 0.357 | 0.235 | 0.802 |
| CMDS(4D)+Ward | 0.451 | 0.342 | 0.262 | 0.451 | 0.568 | 0.357 | 0.235 | 0.802 |
| CMDS(5D)+K-means | 0.571 | 0.224 | 0.261 | 0.565 | 0.727 | 0.396 | 0.248 | 0.787 |
| CMDS(5D)+Ward | 0.571 | 0.268 | 0.453 | 0.556 | 0.727 | 0.357 | 0.319 | 0.787 |
| CMDS(6D)+K-means | 0.565 | 0.248 | 0.504 | 0.392 | 0.722 | 0.521 | 0.288 | 0.752 |
| CMDS(6D)+Ward | 0.495 | 0.495 | 0.504 | 0.634 | 0.727 | 0.660 | 0.220 | 0.752 |
| CMDS(8D)+K-means | 0.392 | 0.300 | 0.294 | 0.392 | 0.717 | 0.521 | 0.255 | 0.773 |
| CMDS(8D)+Ward | 0.357 | 0.248 | 0.248 | 0.580 | 0.727 | 0.521 | 0.314 | 0.773 |
| CMDS(10D)+K-means | 0.502 | 0.300 | 0.294 | 0.555 | 0.717 | 0.357 | 0.255 | 0.777 |
| CMDS(10D)+Ward | 0.502 | 0.380 | 0.248 | 0.599 | 0.798 | 0.521 | 0.314 | 0.766 |
| CMDS(15D)+K-means | 0.357 | 0.308 | 0.357 | 0.555 | 0.722 | 0.357 | 0.308 | 0.777 |
| CMDS(15D)+Ward | 0.502 | 0.320 | 0.404 | 0.599 | 0.798 | 0.521 | 0.261 | 0.766 |
| CMDS(20D)+K-means | 0.412 | 0.249 | 0.357 | 0.392 | 0.798 | 0.484 | 0.261 | 0.766 |
| CMDS(20D)+Ward | 0.357 | 0.248 | 0.248 | 0.503 | 0.717 | 0.521 | 0.248 | 0.787 |
| CMDS(80%)+K-means | 0.309 | 0.275 | 0.275 | 0.565 | 0.389 | 0.248 | 0.268 | 0.773 |
| CMDS(80%)+Ward | 0.309 | 0.248 | 0.255 | 0.556 | 0.439 | 0.289 | 0.220 | 0.773 |
| CMDS(90%)+K-means | 0.392 | 0.300 | 0.504 | 0.555 | 0.727 | 0.357 | 0.248 | 0.737 |
| CMDS(90%)+Ward | 0.357 | 0.380 | 0.504 | 0.599 | 0.727 | 0.521 | 0.319 | 0.787 |
| CMDS(95%)+K-means | 0.357 | 0.300 | 0.357 | 0.555 | 0.773 | 0.456 | 0.255 | 0.766 |
| CMDS(95%)+Ward | 0.502 | 0.380 | 0.248 | 0.599 | 0.809 | 0.521 | 0.314 | 0.787 |
| CMDS(99%)+K-means | 0.369 | 0.249 | 0.357 | 0.392 | 0.798 | 0.484 | 0.261 | 0.766 |
| CMDS(99%)+Ward | 0.357 | 0.248 | 0.248 | 0.599 | 0.717 | 0.521 | 0.248 | 0.787 |

TABLE S9. Leave-one-feature-out clustering ablation for matched JSD distances on the family benchmark. Entries report cluster purity across hierarchical clustering and CMDS-based clustering variants.

| Algorithm | All Features | w/o Structural | w/o Mean Curvature | w/o Hydrogen Bond | w/o Electrostatics | w/o Hydrophobicity | Chemical Only | Geometry Only |
|---|---|---|---|---|---|---|---|---|
| Single | 0.640 | 0.640 | 0.640 | 0.640 | 0.680 | 0.760 | 0.640 | 0.720 |
| Complete | 0.720 | 0.680 | 0.640 | 0.800 | 0.800 | 0.720 | 0.640 | 0.840 |
| Average | 0.680 | 0.640 | 0.640 | 0.800 | 0.800 | 0.720 | 0.640 | 0.840 |
| Weighted | 0.680 | 0.640 | 0.640 | 0.760 | 0.840 | 0.720 | 0.640 | 0.840 |
| CMDS(4D)+K-means | 0.720 | 0.640 | 0.640 | 0.760 | 0.800 | 0.720 | 0.640 | 0.880 |
| CMDS(4D)+Ward | 0.720 | 0.680 | 0.680 | 0.720 | 0.840 | 0.720 | 0.640 | 0.880 |
| CMDS(5D)+K-means | 0.760 | 0.640 | 0.640 | 0.720 | 0.800 | 0.720 | 0.640 | 0.840 |
| CMDS(5D)+Ward | 0.760 | 0.680 | 0.720 | 0.720 | 0.800 | 0.720 | 0.680 | 0.840 |
| CMDS(6D)+K-means | 0.720 | 0.640 | 0.680 | 0.720 | 0.800 | 0.800 | 0.640 | 0.800 |
| CMDS(6D)+Ward | 0.680 | 0.680 | 0.680 | 0.800 | 0.800 | 0.800 | 0.640 | 0.800 |
| CMDS(8D)+K-means | 0.720 | 0.680 | 0.680 | 0.720 | 0.800 | 0.800 | 0.640 | 0.800 |
| CMDS(8D)+Ward | 0.680 | 0.640 | 0.640 | 0.760 | 0.800 | 0.800 | 0.680 | 0.800 |
| CMDS(10D)+K-means | 0.800 | 0.680 | 0.680 | 0.720 | 0.800 | 0.720 | 0.640 | 0.840 |
| CMDS(10D)+Ward | 0.800 | 0.760 | 0.640 | 0.760 | 0.840 | 0.800 | 0.680 | 0.840 |
| CMDS(15D)+K-means | 0.680 | 0.640 | 0.680 | 0.720 | 0.800 | 0.720 | 0.640 | 0.840 |
| CMDS(15D)+Ward | 0.800 | 0.680 | 0.720 | 0.760 | 0.840 | 0.800 | 0.640 | 0.840 |
| CMDS(20D)+K-means | 0.720 | 0.640 | 0.680 | 0.720 | 0.840 | 0.800 | 0.640 | 0.840 |
| CMDS(20D)+Ward | 0.680 | 0.640 | 0.640 | 0.800 | 0.800 | 0.800 | 0.640 | 0.840 |
| CMDS(80%)+K-means | 0.680 | 0.640 | 0.640 | 0.720 | 0.760 | 0.640 | 0.640 | 0.800 |
| CMDS(80%)+Ward | 0.680 | 0.640 | 0.640 | 0.720 | 0.800 | 0.680 | 0.600 | 0.800 |
| CMDS(90%)+K-means | 0.720 | 0.680 | 0.680 | 0.720 | 0.800 | 0.720 | 0.640 | 0.800 |
| CMDS(90%)+Ward | 0.680 | 0.640 | 0.680 | 0.760 | 0.800 | 0.800 | 0.680 | 0.840 |
| CMDS(95%)+K-means | 0.680 | 0.680 | 0.680 | 0.720 | 0.840 | 0.760 | 0.640 | 0.840 |
| CMDS(95%)+Ward | 0.800 | 0.760 | 0.640 | 0.760 | 0.840 | 0.800 | 0.680 | 0.840 |
| CMDS(99%)+K-means | 0.680 | 0.640 | 0.680 | 0.720 | 0.840 | 0.800 | 0.640 | 0.840 |
| CMDS(99%)+Ward | 0.680 | 0.640 | 0.640 | 0.760 | 0.800 | 0.800 | 0.640 | 0.840 |

TABLE S10. Directed threshold success percentages.

| Threshold | Relaxed Top-1 | Relaxed Top-2 | Strict Top-1 | Strict Top-2 |
|---|---|---|---|---|
| 0.30 | 100.00 | 100.00 | 100.00 | 100.00 |
| 0.40 | 100.00 | 100.00 | 98.21 | 100.00 |
| 0.50 | 94.64 | 96.43 | 83.93 | 89.29 |
| 0.60 | 80.36 | 83.93 | 73.21 | 80.36 |
| 0.70 | 75.00 | 78.57 | 66.07 | 69.64 |
| 0.80 | 57.14 | 67.86 | 37.50 | 55.36 |
| 0.90 | 33.93 | 48.21 | 16.07 | 30.36 |

TABLE S11. Bidirectional threshold success percentages.

| Threshold | Strict Top-1 | Strict Top-2 | Relaxed Top-1 | Relaxed Top-2 |
|---|---|---|---|---|
| 0.30 | 100.00 | 100.00 | 100.00 | 100.00 |
| 0.40 | 100.00 | 100.00 | 100.00 | 100.00 |
| 0.50 | 100.00 | 100.00 | 100.00 | 100.00 |
| 0.60 | 78.57 | 92.86 | 96.43 | 100.00 |
| 0.70 | 53.57 | 67.86 | 75.00 | 85.71 |
| 0.80 | 32.14 | 46.43 | 50.00 | 50.00 |
| 0.90 | 7.14 | 21.43 | 25.00 | 42.86 |
